# Adaptive resampling for improved machine learning in imbalanced single-cell datasets

**DOI:** 10.1101/2025.11.04.686583

**Authors:** Zeinab Navidi, Akshaya Thoutam, Madeline Hughes, Srivatsan Raghavan, Peter S. Winter, Lorin Crawford, Ava P. Amini

## Abstract

While machine learning models trained on single-cell transcriptomics data have shown great promise in providing biological insights, existing tools struggle to effectively model underrepresented and out-of-distribution cellular features or states. We present a generalizable Adaptive Resampling (AR) approach that addresses these limitations and enhances single-cell representation learning by resampling data based on its learned latent structure in an online, adaptive manner concurrent with model training. Experiments on gene expression reconstruction, cell type classification, and perturbation response prediction tasks demonstrate that the proposed AR training approach leads to significantly improved downstream performance across datasets and metrics. Additionally, it enhances the quality of learned cellular embeddings compared to standard training methods. Our results suggest that AR may serve as a valuable technique for improving representation learning and predictive performance in single-cell transcriptomic models.

## Introduction

Biological samples contain heterogeneous cellular populations that vary in state and function, and biologically significant properties may arise from rare or underrepresented cell types [1–4]. Understanding this cellular heterogeneity is critical for biological discovery and for diagnostic and therapeutic innovation. Improvements in single-cell RNA sequencing (scRNA-seq) have enabled the study of biology at single-cell resolution [5–7] and the creation of large-scale tissue [8], disease [9], and perturbational [10] atlases. In turn, this has inspired the development of data-driven machine learning (ML) algorithms that learn from these scRNA-seq datasets, with the promise of accelerating downstream biological applications, such as perturbation response prediction, cell type classification, and trajectory inference [11–16].

It is well known that the performance of ML algorithms often depends on the quality and diversity of training data [17–22]. These methods can fail to generalize to out-of-distribution (OOD) contexts when samples are sparsely represented during training [23, 24]. This limitation is particularly important in scRNA-seq applications, where high sample heterogeneity is intrinsic to the data, and rare cell populations can drive key biological processes and disease progression [17, 25–27]. Still, studies have consistently shown that the predictive performance of many ML approaches, including large single-cell foundation models, drops considerably for rarer cell types [12, 28–31]. These challenges underscore the need for robust, generalizable approaches that can learn from imbalanced single-cell datasets and perform well across all cell types [32–34].

Existing approaches to address non-uniform distributions of cell types and states in single-cell modeling include manual upsampling of underrepresented sample sets [30], preselecting or subsampling data based on known annotations (such as marker genes) to create a more balanced training corpora [35], and evaluating model performance using predefined balanced scores to account for imbalanced cell groups in batch integration tasks [29]. However, these approaches still fail to account for the biological context and intrinsic properties of cells, typically require extensive preprocessing prior to training, rely on heuristic-based annotations, and introduce selection biases that compromise model training. More advanced ML techniques to mitigate training data limitations have the potential to improve the abilities of single-cell models to learn from imbalanced datasets [20, 36, 37], but these have yet to be applied to single-cell analyses.

We present an adaptive resampling (AR) algorithm that enhances the ability of ML models to learn from imbalanced single-cell datasets, thereby improving both the quality of their learned embeddings and their performance on downstream tasks. Our method adjusts the training cohort in an online and adaptive manner based on a learned latent projection of training samples, which the AR algorithm leverages to identify underrepresented data points. Through experiments on scRNA-seq datasets with systematic and varied cellular imbalances, we demonstrate that the AR algorithm significantly improves model performance compared to the standard training approach. Our experiments demonstrate the superior performance of AR-trained deep learning models on gene expression reconstruction, cell type classification, and generative perturbation response prediction tasks with respect to a comprehensive set of evaluation metrics and the quality of learned latent embeddings. By addressing disparities in sample representation while remaining independent of user-defined thresholds or predefined criteria, the AR algorithm can be integrated with any model that produces a learned latent distribution and maintains broad applicability across diverse datasets and tasks in single-cell analysis.

## Results

### The adaptive resampling algorithm uses learned latent distributions to reweight data during training

In standard training algorithms for machine learning (ML) models, each input sample is seen once by the model per epoch (Fig. 1A). In this work, we leverage an adaptive resampling (AR) algorithm where the training set is adaptively adjusted at the beginning of each epoch to increase exposure to underrepresented samples and improve learning capabilities for downstream tasks [20]. This objective is achieved by projecting all training samples to a latent space at the beginning of each iteration, calcuating a resampling weight for each sample based on the distribution of latent variables, and then using these resampling weights to select a new combination of samples from the original training set, with replacement (Fig. 1B; Methods).

**Figure 1.**
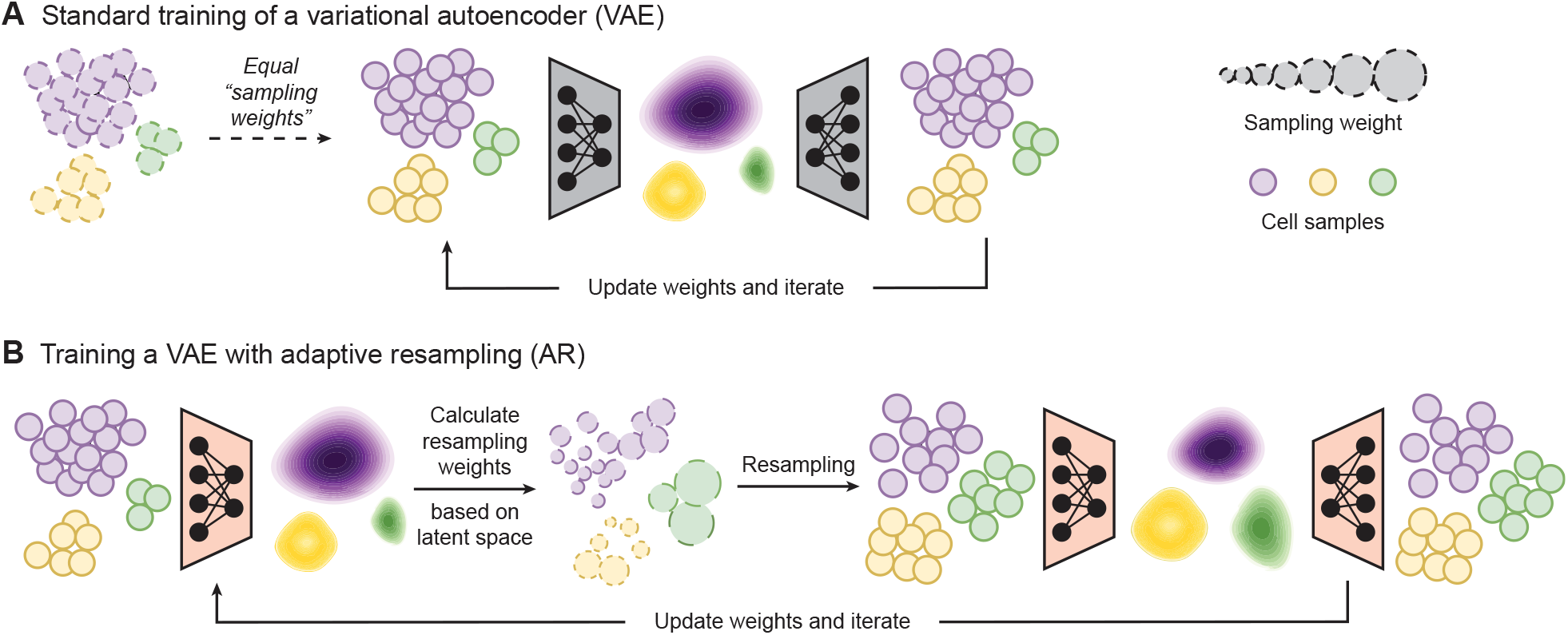
Adaptive resampling for improved cellular representation learning in single-cell datasets. **(A)** Schematic overview of the standard training procedure using a variational autoencoder (VAE) framework as an example. In standard training algorithms, each input sample is seen once by the model per epoch, with individual samples having, in effect, equal “sampling weights”. **(B)** Overview of the Adaptive Resampling (AR) approach, as applied to a VAE. With AR, resampling weights are calculated based on training data distributions in the latent space, increasing the model’s exposure to underrepresented samples during subsequent epochs. This is done by using the resampling weights to select a new combination of samples from the original training set, with replacement.

Specifically, at the beginning of each iteration, all original training samples are projected to a learned latent space, and the probability distribution of each latent variable is estimated. The calculated densities for these latent variables are then inverted and normalized to obtain resampling weights (or resampling score). The inversion step ensures that samples in low-density regions (i.e., underrepresented areas of the latent space) receive higher resampling weights, while samples within high-density regions receive lower weights. Importantly, the resampling weights are adaptively adjusted over the course of model training, as the latent space itself is also refined. This adaptive weighting mechanism identifies underrepresented parts of the training distribution in an entirely unsupervised manner, and thus promotes balanced exposure of samples during training. The final resampling probability for each training data point is obtained by taking the maximum value of resampling scores across all latent dimensions.

The key to the AR algorithm is that underrepresented samples are not determined by the identity or size of predefined groups (e.g., cell types). Instead, sample selection is performed in an unsupervised manner, guided by the latent space which is learned during training and provides a semantically-meaningful compression of high dimensional features. As a result, the proposed AR approach accounts for learned features underlying the data distribution (Fig. 1B). This means that the algorithm functions independently of metadata (e.g., cell type labels or perturbation status), which are often unavailable and have been shown to be inconsistently defined across studies [38, 39]. Furthermore, since the AR strategy operates over latent variables, it can be incorporated into any neural network based model that includes an encoder module. In this work, we utilized variational autoencoders (VAEs) to assess the benefits of AR versus standard training. Implementation, theoretical, and pseudocode details are provided in the Methods.

Our analyses included three single-cell tasks: (1) gene expression reconstruction; (2) cell type classification; and (3) perturbation response prediction. We evaluate the impact of AR on these tasks using established VAE-based architectures [40, 41] and highlight its impact on out-of-distribution (OOD) and rare sample settings in particular, as model generalization to unseen data and under-represented cell populations is a desired, but to date elusive, property for single-cell ML models [12, 23, 28, 28–30].

### Adaptive resampling improves reconstruction of single-cell gene expression profiles

We first asked whether the AR algorithm could enhance representation learning of scRNA-seq data. To this end, we used scVI [40], a well-established scRNA-seq representation learning method that has been shown to outperform more complex deep learning approaches in many fundamental tasks [42, 43]. We trained variants of scVI using both the standard training procedure and the AR approach. Here, we kept all other model hyperparameters and data settings fixed such that the only difference driving model performance was the training algorithm itself. To begin, we evaluated performance on gene expression reconstruction, because having the ability to represent and reconstruct cellular states is fundamental to many downstream single-cell applications.

Previous studies have investigated the impacts that training data composition can have on the generalization capabilities of single-cell deep learning models [18, 19]. To account for these factors, we designed experiments to assess the impact of AR on out-of-distribution, rare, and common samples (all unseen by the models) by varying the diversity of cells that were included during model training (Fig. 2A). Taking data from the large-scale scTab study [44], we used blood cells as a base training cohort and supplemented this group with a varying number of atlas cells (derived from 42 different tissues). Specifically, we maintained a fixed total of 100k cells in the training set. We then trained both the AR and standard scVI variants where 0% (0 cells), 0.001% (1 cell), 0.01% (10 cells), 0.1% (100 cells), 1% (1k cells), 10% (10k cells), and 50% (50k cells) of the total 100k training samples were randomly selected from the atlas population and combined with the base blood cell population (Fig. 2A; Table S1).

**Figure 2.**
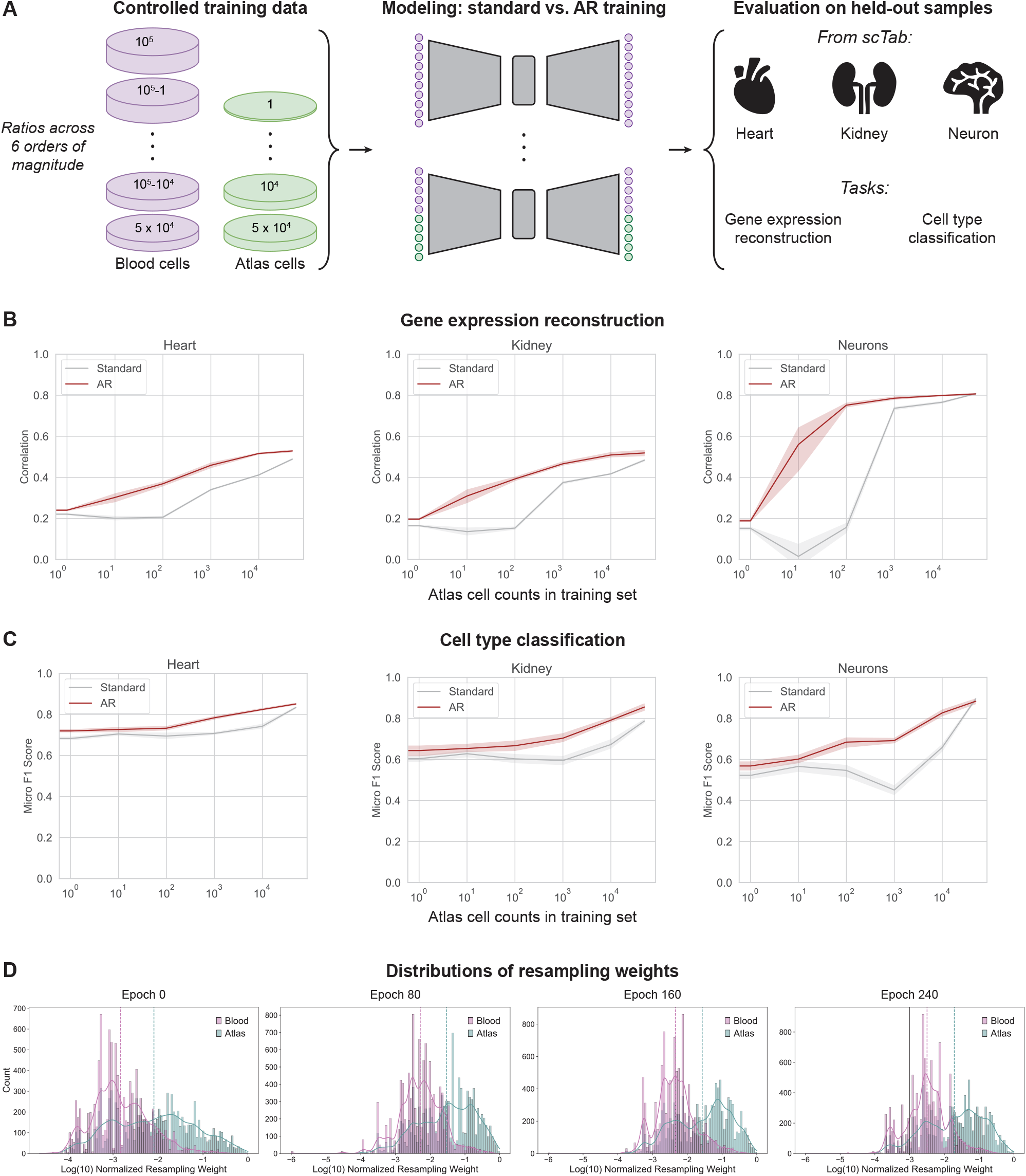
Adaptive resampling enables robust gene expression reconstruction and accurate cell type classification for underrepresented cells. **(A)** Workflow of training data selection, model training, and evaluation on the gene expression reconstruction and cell type classification tasks. **(B)** Gene expression reconstruction performance for scVI models trained in the standard way (grey) or with adaptive resampling (AR; red). Performance is measured by the Pearson correlation coefficient (*r*) over reconstructed gene expression vectors. The x-axis represents the number of atlas cells included in the blood-based training dataset, with all training sets having a total of 100k cells. **(C)** Cell type classification performance from the latent embeddings learned by scVI models trained in the standard way (grey) or with the AR algorithm (red). Performance is measured by the micro F1 Score. The x-axis represents the number of atlas cells included in the blood-based training dataset, with all training sets having a total of 100k cells. Note that results in panels (B) and (C) are based on five different random seeds, with error bars indicating the 95% confidence interval across replicates. **(D)** Distributions of resampling weights for 10k randomly chosen blood training cells and all atlas training cells. The underlying AR-trained scVI model is trained on 90k blood cells and 10k atlas cells. The vertical lines indicate the average of the log_10_-normalized resampling weights for each cell population.

We computed the reconstruction accuracy (via Pearson correlation coefficient, *r*) for each AR and standard-trained scVI model across three evaluation datasets containing heart, kidney, and neuron cells derived from the scTab study (Methods), but different from the pool of atlas cells selected for training (Fig. 2A). Our experiments were designed to simulate predictive scenarios of varying difficulty. Zero-shot evaluations on out-of-distribution samples occured when 0% or 0.001% of atlas cells were included in the training data (i.e., no heart, kidney, and neuron cells were observed during training; Table S1). Scenarios where 0.01%, 0.1%, or 1% of the training data consisted of atlas cells represented cases in which rare cell populations were present during training but at different frequencies. Finally, more balanced evaluations were performed when 10% or 50% of the training data comprised atlas cells. Details on the number of heart, kidney, and neuron cells represented in the training corpus for each analysis are provided in Table S1. Each analysis was repeated across five random seeds for robustness, resulting in a total of 70 experiments.

The AR approach yielded a clear improvement in gene expression reconstruction over the standard algorithm in all training scenarios when evaluating on heart, kidney, and neuronal cells (Fig. 2B). Notably, in this analysis, AR improved model performance with as few as 10 atlas cells seen during training (i.e., 0.01% of the corpus). In constrast, the standard training approach did not benefit from additional atlas cells until at least 1,000 of them were included in the training cohort (i.e., 1% of the corpus)—even then, the standard scVI model never surpassed the performance of the AR-trained version.

As expected, the performance of the AR and standard models converged as more atlas cells were added to the training set, with both approaches achieving approximately the same reconstruction accuracy when 50% of the corpus was atlas cells (Fig. 2B). This observation suggests that there is a point where there is sufficient sample diversity from the atlas cohort to drive performance with standard learning. Moreover, it also suggests that, beyond this point, adding or over-sampling additional atlas cells does not necessarily yield further gains. This is consistent with recent reports on early saturation occuring in single-cell models trained on current large-scale atlases [19]. Together, these findings indicate that the AR algorithm enhances the ability of scVI to represent rare or underrepresented data and improves performance on gene expression reconstuction.

### Adaptive resampling achieves accurate classification of underrepresented cell types in atlas datasets

Next, we investigated if the AR algorithm could enable improved cell type classification, particularly for lowly prevelant cell types observed during training. Here, we evaluated the performance of the trained scVI models, with different training set arrangements of scTab blood and atlas samples (elaborated in the previous section), where model embeddings were used directly with a nearest neighbor classifier to classify cells from the heart, kidney, and neuron evaluation sets (Fig. 2A; Methods).

The AR-trained scVI models significantly outperformed those trained with the standard approach in all three evaluation datasets as measured by micro F1 (Fig. 2C), as well as by accuracy, recall, precision, and macro F1 (Fig. S1; Tables S2-S4). Overall, we observed that the performance of the AR models began to improve noticeably with as few as 100 atlas cells included in the training set; however, the standard approach experienced a delayed performance boost, with relative classification metrics beginning to improve at around 10,000 atlas cells (Figs. 2C and S1). In contrast to the gene expression reconstuction task, the standard models required greater numbers of atlas cells in training to yield performance gains in the cell classification task compared to the setting with no atlas cells included.

To investigate which cells were being preferentially resampled, we assessed the distribution of sample-wise resampling weights over the course of AR model training both quantitatively (Fig. 2D) and qualitatively (Fig. S2). We were able to more clearly observe the resampling weight updates in settings where a large number of atlas cells were included in the training set (e.g., 10,000 cells making up 10% of the corpus). We used uniform manifold approximation and projection (UMAP) to visualize the landscape of training samples annotated both by their source group (blood versus other atlas cells) and by their normalized resampling weights across four epochs at different stages of AR model training (Fig. S2). We also investigated the total distribution of resampling weights across blood and atlas training cells (Fig. 2D). Here, our results showed that atlas cells had overall higher resampling scores compared to blood samples, suggesting that the AR algorithm recognized these cells as being underrepresented based on their projected latent distribution, ultimately leading to enhanced predictive performance. These findings highlight the ability of the AR algorithm to encourage models to identify and preferentially sample relatively rarer samples during training.

### Adaptive resampling enables generalization in perturbation response prediction

We next asked whether the adaptive resampling approach could confer benefits on more complex scRNA-seq tasks. To this end, we considered single-cell perturbation response prediction, which aims to predict the gene expression profile resulting from a specific perturbation, given the unperturbed gene expression profile as input. This task is key to understanding how gene expression changes in response to cellular signaling cues, such as cytokines, or pharmacological interventions, such as treatment with small molecules drugs. We focused on perturbation response in particular given the importance of OOD generalization to this task, as its downstream biological utility relies on *in silico* prediction of perturbation effects in experimentally unseen or unmeasured cellular contexts.

We hypothesized that the AR algorithm could improve the generalization of cellular perturbation response prediction to held-out cell groups. To test this, we used scGen [41] as the base VAE model, with each of the experiments described below run across five independent trials. In each experiment, we defined each annotated cellular subpopulation as a separate cell group (either species-based or cell type-based, depending on the datasets) and excluded perturbed cells from one cell group (considered as the OOD test group) during model training. The objective during inference was to predict the counterfactual perturbed gene expression profiles for cells from the held-out cell group, evaluated by comparing model predictions against the ground truth perturbed gene expression measurements (Fig. 3A; Methods).

**Figure 3.**
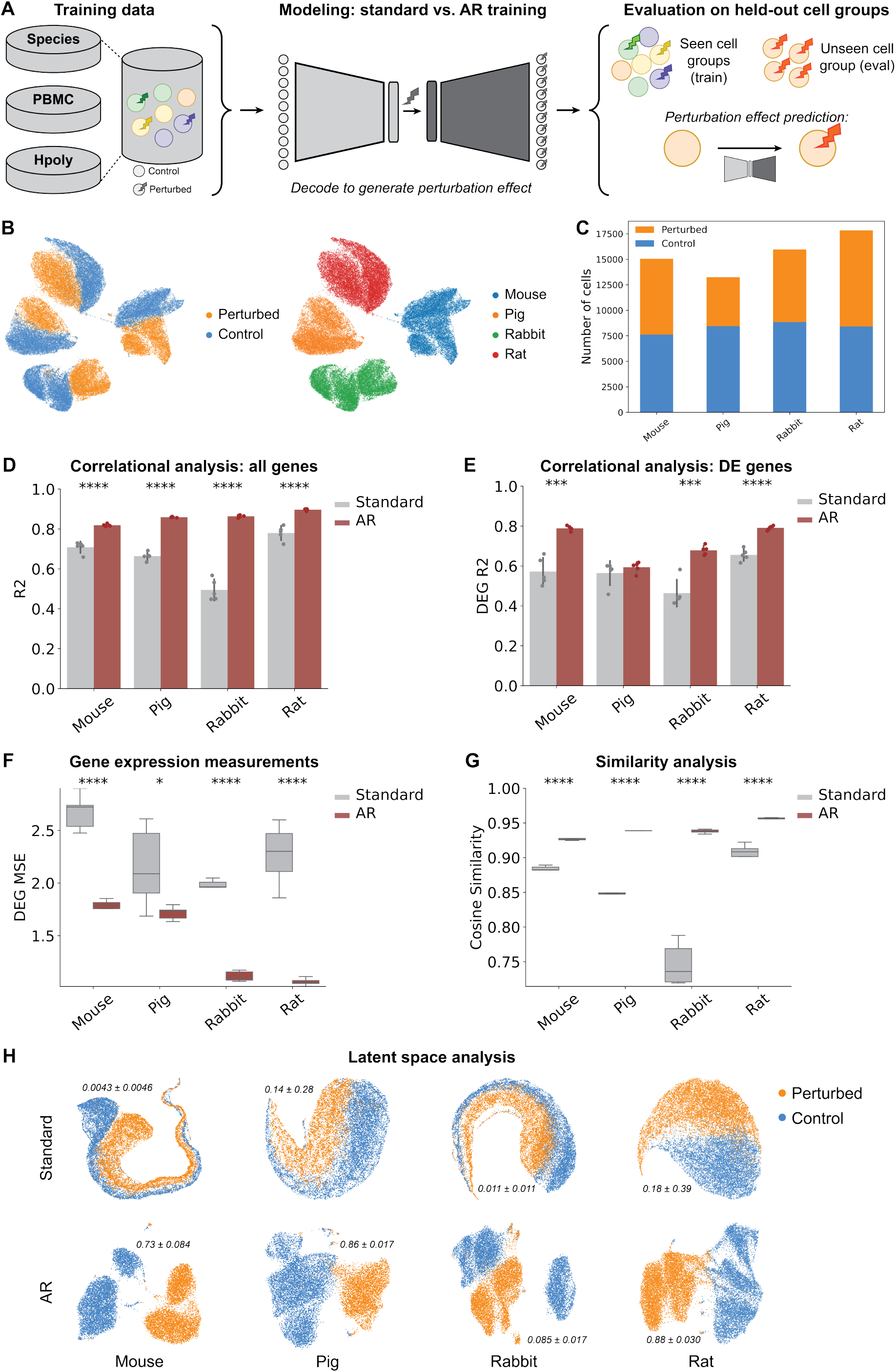
Adaptive resampling enables robust prediction of immune cell stimulation across species. **(A)** Workflow of training data selection, model training, and evaluation on the single-cell perturbation response prediction task. **(B)** UMAP visualization of sample distributions in the Species dataset, annotated by condition (control versus perturbed with LPS, left) and by cell groups (individual species, right). **(C)** Cell counts per condition and species. **(D-E)** Square of the Pearson correlation coefficient between real and predicted perturbed gene expression using AR versus standard models for each held-out test cell group across (D) all genes and (E) only differentially expressed genes. Results are based on five different random seeds, with error bars representing the standard deviations across replicates. **(F)** Mean sqaured error (MSE) computed between ground-truth and predicted perturbed gene expression profiles across differentially expressed genes for AR versus standard models. **(G)** Distributions of cosine similarity scores for predicted expression profiles across all genes, relative to ground-truth expression profiles. **(H)** UMAP visualizations of the latent space projections for test samples, annotated by condition, comparing AR and standard learned latent embeddings. The adjusted Rand index (ARI) is reported; the ARI evaluates the clustering performance of a K-means algorithm applied to the embeddings derived from each model, comparing the predicted clusters to ground truth labels of control and perturbed cells. Two-sided t-tests were performed in all experiments to compare the means of metrics, with ^*^*P* < 0.05, ^**^*P* < 0.01, ^***^*P* < 0.001, and ^****^*P* < 0.0001.

We evaluated our hypothesis on three public scRNA-seq datasets representing different perturbation and generalization scenarios. We refer to these as the Species [45], PBMC [46], and H. poly [47] datasets. The Species dataset is composed of bone marrow derived mononuclear phagocytes derived from mice, rats, rabbits, and pigs that were stimulated with lipopolysaccharide (LPS) [45]. The PBMC dataset is composed of seven different cellular subpopulations of human peripheral blood mononuclear cells (PBMCs) captured either in a baseline control state or after being stimulated with interferon (IFN-*β*) [46]. The H. poly dataset measured the responses of murine epithelial cells, annotated into eight different subpopulations, to pathogen infection from the parasitic helminth *Heligmosomoides polygyrus* [47]. Each cell group within each dataset contained both control and perturbed samples. We observed the most separated distribution of cell groups in the Species dataset (Fig. 3B) and characterized the split of control-versus-perturbed cells across these cell groups (Fig. 3C). Because the Species dataset represents a scenario with the most disparate populations and data sources for which we aim to generalize, we focus on results for the Species dataset in the main text and present complete results for the PBMC and H. poly datasets in Figures S3 and S4.

We measured the correlation between ground truth and predicted perturbed gene expression profiles via the squared Pearson correlation coefficient (*R*^2^) over all genes (Figs. 3D, S3C, and S4C) and differentially expressed genes (DEGs) between perturbed and control cells (Figs. 3E, S3D, and S4D). On the Species dataset, the incorporation of AR significantly improved perturbation response prediction on cells from the held-out species. The magnitude of this performance boost differed across cell groups, where in some cases the *R*^2^ for AR models nearly doubled relative to that for standard training (e.g., on rabbit cells). We further evaluated model performance by comparing predicted and ground truth gene expression profiles using mean squared error (MSE) and cosine similarity considering all genes and DEGs for different held-out cell groups. The MSE was significantly reduced for AR-trained models across all datasets with varying biological characteristics (Figs. 3F, S3E, and S4E), and AR-trained models yielded higher cosine similarity between predicted and ground truth gene expression profiles (Figs. 3G, S3F, and S4F). Moreover, across all metrics, the AR models exhibited more consistent performance with smaller variance across independent trials, suggesting a degree of conferred stabilization and regularization.

To measure how well AR enabled generalization in the presence of rare samples, we trained a suite of scGen models on all cell groups except one, withholding both control and perturbed cells from that group. We then trained models where we incrementally introduced varying proportions of control cells from the held-out group into the training corpus—specifically, 0%, 20%, 40%, 60%, 80%, and 100%. During inference, all control cells from the test group were provided as individual inputs, and predictions were compared against the true expression profiles of perturbed cells. These control sets, depending on their inclusion level, represented completely out-of-distribution (i.e., 0%), rare, or common sample distributions encountered by the model during training. We observed that using AR led to better model performance in all configurations, and that it benefitted from having seen as little as 20% of the control cells, while standard training did not (Figs. S5-S7). Altogether, these findings suggest that the AR algorithm can learn more effectively from rare cellular patterns than standard training procedures to improve overall performance and generalizaltion capabilities in perturbation response prediction.

Finally, we asked whether these performance gains stemmed from a more informative latent space learned by AR-trained models compared to those trained with the standard algorithm. Here, we observed that the AR-derived embeddings of test samples exhibited qualitatively less overlap and more regularized distributions between control and perturbed cells, whereas embeddings from standard-trained models showed greater mixing of these conditions (Figs. 3H, S3G, and S4G).

We quantitatively evaluated how well the learned latent representation preserved cellular conditions (control versus perturbed). Specifically, we computed the adjusted Rand index (ARI) by performing K-means clustering on the latent embeddings and comparing the resulting clusters with the ground-truth labels (Fig. 3H). Overall, the ARI scores indicate that the embeddings generated by our AR approach preserve biologically relevant information more effectively than those produced by the standard model. Learning structured and discriminative embeddings is particularly valuable, as these representations can be repurposed for diverse downstream bioinformatic analyses. Notably, the UMAP projections of the latent embeddings for the Species dataset had more pronounced differences between AR versus standard scGen models than those generated for PBMC and H. poly datasets. We attribute this to the fact that the PBMC and H. poly cell groups are qualitatively more similar (Figs. S3A, S4A), whereas the sample groups in the Species dataset are distributionally distinct (Fig. 3B). These findings suggest that the use of the AR approach can provide the greatest benefit when observations exhibit strong intrinsic heterogeneity, yielding latent spaces that are comparable (PBMC and H. poly) and sometimes superior (Species) to those obtained with standard training.

## Discussion

Single-cell datasets inherently have non-uniform distributions of diverse cell types and states. Thus, machine learning algorithms trained on these datasets inherently learn patterns from the more abundant cell populations, leading to poor model performance for unobserved and rarer samples [12, 23, 24, 28, 28–30]. Here we present an Adaptive Resampling (AR) algorithm to mitigate biases introduced by imbalanced cellular representation in computational modeling of single-cell RNA-seq datasets. We demonstrate the efficacy of AR in improving the performance and generalization of variational autoencoder based scRNA-seq analysis tools through evaluation on gene expression reconstruction, cell type classification, and perturbation response prediction on unseen out-of-distribution, underrepresented, and more common sample sets. Our results show that incorporating AR as a density-aware resampling technique into existing generative models for scRNA-seq yields significant improvements in downstream prediction performance, stabilizes results, and enhances the quality and richness of latent cell embeddings.

The adaptive resampling algorithm utilizes a learned distribution over latent variables to define resampling weights for each training sample. This is done in a fully unsupervised manner by projecting samples into a latent space and assigning higher resampling weights to observations that have lower probabilities. The resampling weights are updated adaptively during training, and thus AR exposes underrepresented samples to the model with higher frequency, alleviating unwanted bias or overfitting towards more abundant cell groups.

Our experiments demonstrate that AR improves model generalization to unseen sample sets and rare cellular populations across different tasks and applications. When reconstructing gene expression profiles, AR-trained models yield significant improvements, suggesting higher quality learned cell embeddings. This is further supported by significant enhancements in downstream performance in cell type classification conducted directly from these cell embeddings. Finally, our evaluations showing that AR significantly improves perturbation response prediction are particularly noteworthy, as they suggest that adaptive resampling has the potential to strengthen machine learning capabilities for predicting perturbations in rare and unseen cell groups, an outstanding current challenge in the field [32, 48, 49]. Because it utilizes a learned latent variable distribution, the AR algorithm does not require any sample selection of cells prior to training nor any metadata, which can be costly or even infeasible to collect in many cases. As a result, it can be easily integrated into any single-cell modeling framework that includes an encoding module.

While our evaluations span three different scRNA-seq tasks, AR being implemented within two different models, and analyses on eight different datasets, our work is not without limitations. For example, all our experiments were constrained to VAE-based architectures. Even though we made this choice to validate the impact of AR in performant architectures that directly learn a latent space, as we mentioned, the AR technique could feasibly be incorporated into any neural network-based model that includes an encoder component. Investigating the influence of the AR algorithm when combined with non-VAE models, such as transformer-based encoders [50], is a focus for future work. Relatedly, the AR algorithm can be applied in many different ways. In this work, resampling occurs at the beginning of each epoch to adjust which observations are used during training. Alternatively, resampling weights could be computed at the start of training using an already frozen or pretrained model, and the training samples then resampled accordingly. This latter scenario is particularly relevant in the context of large-scale single-cell foundation models, where developers can leverage foundation model embeddings to define training sample weights and perform resampling prior to training a new ML framework for a desired task. We provide a modular implementation of the AR algorithm in our GitHub repository to calculate resampling weights for either setting (see Methods). Another interesting direction for future work would be to combine the AR algorithm with previous supervised approaches also aimed to address data imbalances in biological applications. Strategies like upsampling enhance training set diversity based on cell type labels, but AR captures patterns that are latent and go beyond *a priori* annotations, which are often inconsistently defined across studies. Pairing these two strategies together could lead to even better performance gains for single-cell models. Lastly, with the advent of atlases containing tens to hundreds of millions of cells with various multi-omic measurements [10, 51], there is great opportunity to extend the AR framework to the multimodal setting.

In sum, our work presents adaptive resampling as an effective approach to enhance the learning and generalizability of ML models for the analysis of heterogenous scRNA-seq datasets. The AR algorithm confers the ability to learn more informative and better-regularized cell embeddings, which can be independently utilized for diverse downstream analysis. Given the emergence of single-cell foundation models that attempt to learn high-quality cell embeddings, we envision that the AR algorithm will be particularly relevant to enhancing the quality and utility of these embeddings across a broad range of tasks and contexts. We hope that this work inspires future research into better utilization of the rich heterogeneity of cellular representation in single-cell datasets for both biological discovery and therapeutic applications.

## Author contributions

Conceptualization: Z.N., L.C., A.P.A.; Methodology: Z.N., L.C., A.P.A.; Software Programming: Z.N., A.T., M.H., A.P.A.; Experimental Design: Z.N., L.C., A.P.A.; Investigation: Z.N., A.T., M.H., L.C., A.P.A.; Validation: Z.N., A.T., M.H., L.C., A.P.A.; Formal analysis: Z.N., A.T., M.H., L.C., A.P.A.; Resources Provision: L.C., A.P.A.; Visualization: Z.N., A.P.A.; Writing - Original Draft: Z.N., A.T., L.C., A.P.A.; Writing - Review & Editing: Z.N., A.T., M.H., S.R., P.S.W., L.C., A.P.A.; Supervision: S.R., P.S.W., L.C., A.P.A.

## Acknowledgements

The authors thank Nicolo Fusi for project feedback, Sean Whitzell for assistance with compute resources, and Stephanie Simmons and Hannah Richardson for assistance with software release. Z.N. greatlfully acknowledges the support of her PhD supervisors, Dr. Bo Wang and Dr. Benjamin Haibe-Kains, during the completion of this project. S.R. acknowledges funding support from NCI K08 CA260442. Any opinions, findings, and conclusions or recommendations expressed in this material are those of the author(s) and do not necessarily reflect the views of any of the funders.

## Methods

### Overview of the adaptive resampling algorithm

Consider a dataset **X** ∈ *R*^*N*×*D*^ with *n* = 1, …, *N* training samples, where each sample has *d* = 1, …, *D* features. We will use **x**_*n*_ = [*x*_*n*1_, …, *x*_*nD*_] to denote the row vector of **X** (i.e., the gene expression of the *n*-th cell). Assume **X** is composed of *M* distinct subgroups (e.g., cell types, cell lines, or species) with an imbalanced number of samples per group. The core idea of the AR algorithm is to define a resampling weight for each *i*-th sample and use it as the probability of selecting that sample during each training iteration, with replacement. These weights are designed to downweight overrepresented samples that lie in dense regions of the latent space and upweight rarer samples, encouraging the model to focus on underrepresented data points.

At the beginning of each iteration, resampling weights of all training samples are initialized to zero. The encoder component maps each observation in **X** to a *K*× *N* lower dimensional space characterized by **Z**, where each **z**_*k*_ (where *k* = 1, …, *K* and *K* ≤ *D*) is assumed to approximately follow some distribution *q*_*ϕ*_(**z** |**X**) parameterized by parameters *ϕ*. For each *k*-th latent dimension the projected values across all *N* samples are used to estimate a probability distribution function *p*(**z**_*k*_). Next, we smooth over this probability distribution function and normalize across all observations to obtain a per-sample probability where

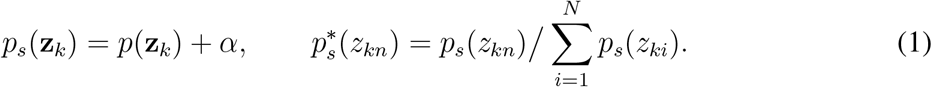

Here, the smoothing factor *α* is a fixed hyperparameter used to tune the degree of debiasing [20]. To obtain unnormalized resampling weights, we invert each per-sample probability such that

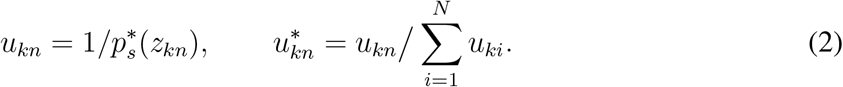

The inversion step ensures that samples falling into low-density regions (i.e., underrepresented areas of the latent space) receive higher resampling weights, while samples within high-density regions receive lower weights. This adaptive weighting mechanism promotes balanced exposure of samples during training, improving representation learning in imbalanced settings. The above calculation is repeated for each individual *k*-th latent dimension, and the final unnormalized resampling weight for the *n*-th sample is computed as: 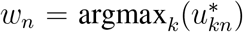. This step ensures that the algorithm focuses on samples that are underrepresented in at least one latent space dimension, regardless of their global rarity. As a last step in the algorithm, the resampling weights are normalized so that they represent the probability of resampling each training data point

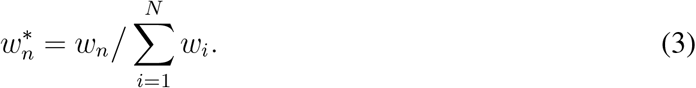

The (normalized) resampling weights in Eq. (3) represent the probability of sampling a given cell (with replacement) to be included in the *N* -sized training set at the beginning of the next iteration. We will denote this resampled data as **X**^*^, which will be fed into the model where we calculate the loss between the predicted values (*ŷ*) and ground truth (*y*), followed by optimization of model parameters. This is different from the standard training approach where all training samples are fed to the model once.

Once again, the AR algorithm is compatible with a wide rage of neural network architectures that include an encoder module. The encoder can be intrinsic to the model (i.e., as in a variational autoencoder) or implemented as a component within a larger modeling framework (i.e., as in the architecture proposed in Amini et al. [20]). In the latter case, the overall pipeline can jointly perform two tasks: (1) optimizing the original predictive objective and (2) learning the latent representation of samples via a reconstruction loss. Pseudocode illustrating the general steps of the AR framework can be found in Algorithm 1. The code for the AR algorithm used in this study was inspired by a prior study in computer vision from Amini et al. [20], as well as by a corresponding implementation of the framework from Kang [52].

#### Algorithm 1

Adaptive Resampling (AR) Algorithm

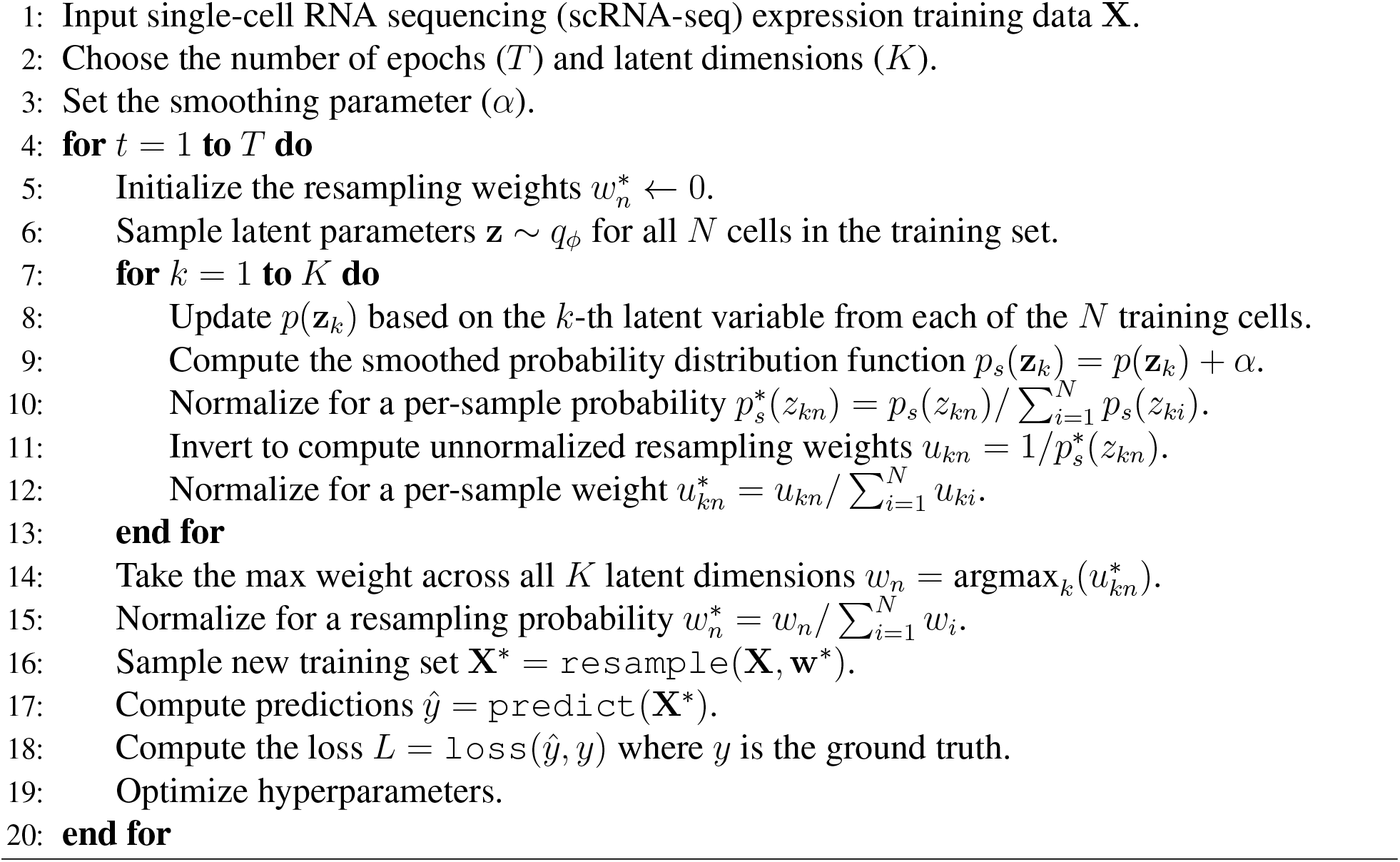

#### Hyperparameter settings for the AR algorithm

There are two hyperparameters used in the implementation of the AR algorithm. The first is the smoothing factor *α*, which was set to 0.0001 in our experiments. The second parameter determines the number of bins used to estimate the probability distribution for the latent variables which we do by calling the `numpy.histogram()` function. While our experiments did not reveal a strong sensitivity to small changes in these parameters, they may be adjusted depending on the number of latent dimensions and the range of projected values.

#### Implementation details

We integrated the AR algorithm into the original training algorithm for each single-cell analysis tool by modifying their publically available source code or using their application program interface (API). Apart from resampling the training cohort fed into the model at each iterataion, all other components of the algorithm, along with all experimental settings, were held constant between the AR and standard training procedures across all experiments. This setup enabled us to isolate and evaluate the direct impact of the adaptive resapmling strategy on downstream performance.

### scVI

scVI (single-cell variational inference) by Lopez et al. [40] is a well-known deep generative model for scRNA-seq applications and is implemented as a variational autoencoder (VAE). It is designed to support downstream predictive tasks such as cell type annotation and gene expression reconstruction. The primary objective of scVI training is to generate a denoised and biologically informative latent space from single-cell transcriptomic profiles while preserving the model’s ability to reconstruct the original expression data.

Briefly, the model uses variational inference to approximate an intractable posterior distribution for the latent variables *p*(**z** | **X**) with a variational approximating family *q*_*ϕ*_(**z** | **X**) parameterized by *ϕ*. The training objective in this framework is to maximize an “evidence lower bound” (ELBO). When training scVI, we extracted **z** ∼ *q*_*ϕ*_ for each training sample to compute the resampling weights using the `get_latent_representation()` function as part of the `scvi-tools` package (release 1.0.4) [53]. Again, the model architecture was left unchanged, with the latent dimensionality set to *K* = 64. All other architectural details, such as dropout and layer size, were left at their default settings. We trained each model without early stopping for 300 epochs while using a batch size of 4096 and a learning rate of 5× 10^−5^ on a NVIDIA RTX A6000. We disabled early stopping to keep the training procedure consistent across each of the trained models, and the final model was selected based on the best validation loss.

### scGen

scGen by Lotfollahi et al. [41] combines variational autoencoders and latent space vector arithmetic for predicting cellular responses to perturbations. An example of this task is inferring the gene expression profile of drug-treated cells using data from untreated cells. In practice, scGen is trained on both control and perturbed cells where the aim is to learn a latent space that captures differences between conditions. The average shift between the states in the latent space is stored as a vector ***δ***.

During inference, the latent representation of a control cell within the held-out test set is shifted by *δ* to estimate the latent representation of the perturbed cellular state: **z**_*p*_ = **z**_*c*_ + ***δ***. The decoder then projects **z**_*p*_ back into the high-dimensional gene expression space. While implementing the AR strategy during model training, the latent embeddings were extracted using the `SCGENVAE.inference()` function and then used to compute the resampling weights.

In this work, we used the official scGen package for our analyses (release 2.1.1). The model architecture had *K* = 64-dimensional latent space, the batch size was set to 2048, and both the learning rate and weight decay were set to 5× 10^−5^. Each model was trained for a fixed number of epochs: 1000 for the Species datasets, 1800 for the PBMC datasets, and 2000 for the H. poly datasets. Once again, early stopping was disabled to keep the training procedure consistent across models, and the final model was selected based on validation loss. All training and validation experiments were performed using a NVIDIA RTX A6000.

### Datasets for gene expression reconstruction and cell type classification

The main dataset used for training and evaluation on the gene expression reconstruction and cell type classification tasks was scTab [44], which is a subset of the CELLxGENE census [54] (version 2023-05-15). scTab consists of 22.2 million cells from 164 unique cell types, 5,052 unique human donors, and 56 human tissues.

#### Pretraining datasets

We constructed two datasets from scTab (also see Nadig et al. [18]).

- **Blood cells (base):** This dataset was created by subsampling exclusively from the blood cell populations within scTab.
- **Atlas cells (spiked-in):** This dataset was created by taking cells from scTab regardless of tissue label.

For preprocessing, we filtered out cells with a proportion mitochondrial counts greater than 0.1 and a total number of unique molecular identifiers (UMIs) greater than 1000. We pretrained scVI using seven distinct combinations of the blood and atlas datasets, each comprising 100k cells. These combinations were designed to simulate varying degrees of cell type imbalance in the training corpus of a deep learning model. Specifically, 0%, 0.001%, 0.01%, 0.1%, 1%, 10%, and 50% of the total 100k training samples were randomly selected from the atlas population and combined with the base blood cell population. These mixtures included: (1) 100,000 blood cells + 0 atlas cells, (2) 99,999 blood cells + 1 atlas cell, (3) 99,990 blood cells + 10 atlas cells, (4) 99,900 blood cells + 100 atlas cells, (5) 99,000 blood cells + 1000 atlas cells, (6) 90,000 blood cells + 10,000 atlas cells, (7) 50,000 blood cells + 50,000 atlas cells. For each case, the final 100k cells were divided into 80/20 splits for training and validation. Note that all atlas cells were uniformly downsampled from scTab (i.e., sampled at random without any consideration of additional metadata or expression counts).

#### Evaluation datasets

We evaluated the standard and AR trained scVI models on three datasets, with these evaluation samples being different from the samples seen by the models during training.

- **Heart**. This evaluation dataset was created by subsampling all the cells labeled “heart” in scTab. Here, we retained only those with the following cell type labels: “cardiac muscle cell”; “pericyte”; “capillary endothelial cell”; “fibroblast of cardiac tissue”; “endothelial cell of artery”; “smooth muscle cell”; “endothelial cell”; “vein endothelial cell”; “neuron”; “endothelial cell of lymphatic vessel”; “cardiac neuron”; and “mesothelial cell”. The final evaluation dataset contained 13,571 cells.
- **Kidney**. This evaluation dataset was created by subsampling all the cells labeled “kidney” in scTab. Here, we retained only those with the following cell type labels: “epithelial cell of proximal tubule”; “kidney loop of Henle thick ascending limb epithelial cell”; “endothelial cell”; “kidney collecting duct principal cell”; “kidney distal convoluted tubule epithelial cell”; “kidney interstitial fibroblast”; “kidney collecting duct intercalated cell”; “kidney connecting tubule epithelial cell”; “kidney loop of Henle thin descending limb epithelial cell”; “kidney loop of Henle thin ascending limb epithelial cell”; “renal interstitial pericyte”; and “vascular associated smooth muscle cell”. The final evaluation dataset contained 8,468 cells.
- **Neuron**. This evaluation dataset consisted of a random set of 10,000 neuronal cells subsampled from the Human Brain Cell Atlas (HBCA) [55]. The HBCA resource is publicly available online and can be accessed via GitHub at github.com/linnarsson-lab/a dult-human-brain.

For each of these datasets, we performed the same filtering for mitochondrial read proportion and number of UMIs as described above for the training datasets. The numbers of cells listed above describe the post quality-controlled sample sizes per dataset.

### Datasets for perturbation response prediction

To investigate the effect of the AR training strategy on perturbation response prediction, we downloaded the following preprocessed datasets from Lotfollahi et al. [41]. Note that each contains control and perturbed samples provided for all cell groups:

- **Species**. This dataset from Hagai et al. [45] is composed of bone marrow-derived mononuclear phagocytes from mice, rats, rabbits, and pigs stimulated with lipopolysaccharide (LPS) for six hours. It contains 62,114 cells in total with 6,619 highly variable genes which were used for our analyses.
- **PBMC**. This dataset from Kang et al. [46] includes two groups of human peripheral blood mononuclear cells (PBMCs) in both a control state and stimulated with interferon (INF-*β*). In total, there are 16,893 cells representing 8 different cell types, and 6,998 highly variable genes were selected for our analyses.
- **H. poly**. This dataset from Haber et al. [47] contains epithelial cell responses to pathogen infection where the responses of murine intestinal epithelial cells to *Salmonella* and parasitic helminth *H. poly* were investigated. We focus on the *H. poly* subset. In total, there were 5,059 cells, with the top 7,000 highly variable genes being used for our analysis.

For each dataset, the read counts were normalized per cell, and gene expression was log-transformed to facilitate a smoother training procedure with scGen Lotfollahi et al. [41].

### Evaluation setup and metrics

We compared the utility of training deep learning models with the AR approach on three different downstream tasks: gene expression reconstruction, cell type classification, and perturbation response prediction. The specific evaluation strategies for each are detailed below.

#### Gene expression reconstruction

For each single-cell sample, we compared the log-transformed and normalized gene expression counts from the original data with the reconstructed profiles generated by scVI (total library size of 10,000). The Pearson correlation coefficient between real and predicted normalized gene expression profiles was calculated and averaged across all cells in the evaluation datasets (again Heart, Kidney, and Neuron).

#### Cell type classification

For each evaluation dataset (i.e., Heart, Kidney, and Neuron), gene expresion profiles were first projected to latent embeddings that were generated using the trained scVI models. The resulting emeddings were then randomly split into two equal halves. We trained a nearest neighbors classifier (with *k* = 5) on the first half (using `KNeighborsClassifier()` from sklearn package), and predictions were made on the second half. Performance was evaluated by comparing these predictions to the ground truth annotations. Evaluation metrics included accuracy, precision, recall, micro F1, and macro F1 scores.

#### Perturbation response prediction

For each held-out test cell group (i.e., where the perturbed gene expression profiles were excluded from scGen during model training), we compared the average predicted expression of individual genes, averaged over all model predictions (i.e., input control cells), to the average ground-truth expression, averaged over all real perturbed cells. We employed different complementary evaluation metrics for a holistic view of downstream performance.

- **Squared Pearson correlation coefficient (*R***^***2***^). This metric assesses how well the model’s predictions capture the overall regulation of gene expression, relative to the ground truth expression profile.
- **Mean squared error (MSE)**. This metric measures the absolute difference between predicted and actual gene expression values for the perturbed cells.
- **Cosine similarity**. This metric evaluates the similarity between predicted and real gene expression vectors in high-dimensional space, reflecting representational similarity.
- **Latent space analysis**. These visualizations qualitatively assess how the AR algorithm influences the structure and regularization of the learned cell embeddings compared to standard training. The adjusted Rand index (ARI) was used to assess the extent to which the biological differences between control and perturbed cells were preserved in the latent embeddings of test samples produced by the AR and standard scGen models.

In the perturbation response prediction tasks, we also designed an additional evaluation to investigate how varying the inclusion of control cells from the held-out test group would affect model performance. Here, we randomly included 0%, 20%, 40%, 60%, 80%, and 100% of these control cells from the test group during training, while all perturbed test cells were consistently excluded. The task during model training is to reconstruct the input gene expression profile. During inference, all control cells from the test group were used as input, identical across all configurations, and the model’s output was the predicted perturbed gene expressions. Importantly, this setting aimed to evaluate how the AR and standard training procedures would perform under varying degrees of exposure to samples with similar distributions to the test set (see Figures S5-S7).

#### Statistical analysis

In all the evaluations detailed above, we perfomed two-sided t-tests to assess statistical significance between the AR and standard trained models. In figures, significance levels are indicated in the plots as follows: ^*^*P* < 0.05, ^**^*P* < 0.01, ^***^*P* < 0.001, and ^****^*P* < 0.0001.

### Code and data availability

All code is available under a permissive MIT license at https://github.com/microsoft/sc-AR. All preprocessed data files used for the analyses conducted in this paper can be downloaded from the project’s Zenodo dashboard at https://zenodo.org/records/15186018.

Here, we utilized scVI [40] which is implemented as part of scvi-tools [53] and downloaded from https://scvi-tools.org/ (release 1.1.2). We also used scGen [41] (release 2.1.1) which can be downloaded from https://github.com/theislab/scgen.

The blood cells, atlas cells, heart, kidney, and neuron datasets are all available in the CELLxGENE census and can be found online at https://cellxgene.cziscience.com/. Instructions for downloading the scTab corpus from the CELLxGENE census are provided at https://github.com/microsoft/scFM-dataselection/tree/main/data/preprocess. The preprocessed Species, PBMC, and H. poly datasets used in this study are directly downloaded from Lotfollahi et al. [41] at https://drive.google.com/drive/folders/1v3qySFECxtqWLRhRTSbfQDFqdUCAXql3.

## Supplementary Figures and Tables

**Table S1.** Mean and standard deviation of Heart, Kidney, and Neuron cell counts included during scVI model training for five different seeds across different scenarios where 0% (0 cells), 0.001% (1 cell), 0.01% (10 cells), 0.1% (100 cells), 1% (1k cells), 10% (10k cells), and 50% (50k cells) of the total 100k training samples were randomly selected from the atlas population and combined with the base blood cell population.

| # Atlas Cells | 0 | 1 | 10 | 100 | 1000 | 10000 | 50000 |
| --- | --- | --- | --- | --- | --- | --- | --- |
| Heart | 0.0 ± 0.0 | 0.0 ± 0.0 | 0.4 ± 0.5 | 2.6 ± 1.5 | 19.8 ± 1.5 | 179.4 ± 7.8 | 928.2 ± 31.3 |
| Kidney | 0.0 ± 0.0 | 0.0 ± 0.0 | 0.2 ± 0.4 | 0.8 ± 0.8 | 12.2 ± 2.6 | 122.2 ± 10.4 | 593.8 ± 9.9 |
| Neuron | 0.0 ± 0.0 | 0.0 ± 0.0 | 1.8 ± 0.4 | 23.4 ± 3.1 | 224.6 ± 7.1 | 2221.2 ± 20.7 | 11256.8 ± 34.2 |

**Table S2.**
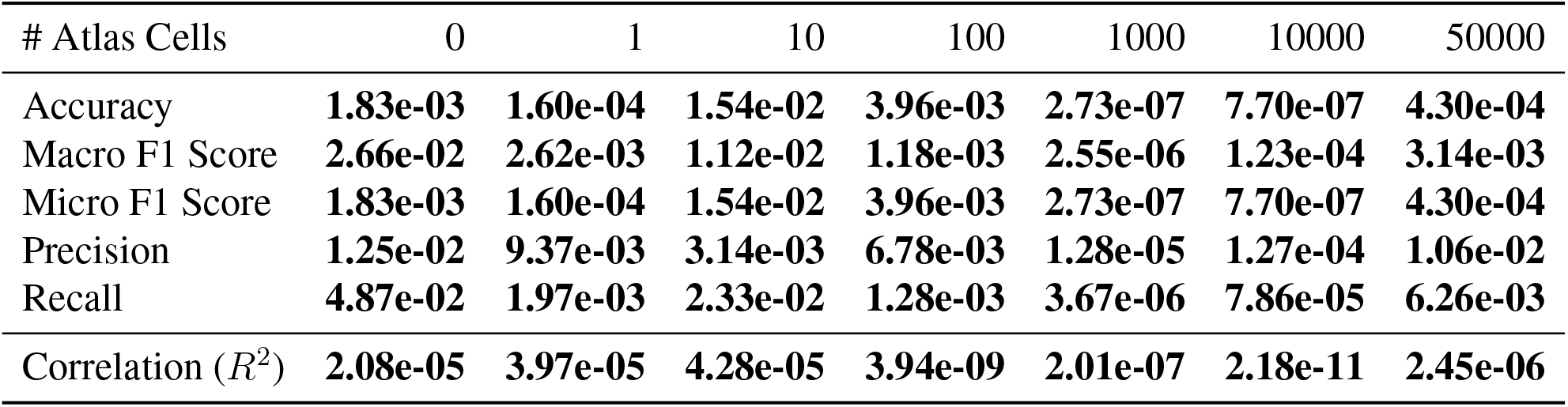
The p-values for two-sided t-tests on evaluation metrics for the AR and standard scVI models tested on the Heart dataset. Bold indicates statistical significance as determined by *P* < 0.05. Results are based on five replicates.

| # Atlas Cells | 0 | 1 | 10 | 100 | 1000 | 10000 | 50000 |
| --- | --- | --- | --- | --- | --- | --- | --- |
| Accuracy | **1.83e-03** | **1.60e-04** | **1.54e-02** | **3.96e-03** | **2.73e-07** | **7.70e-07** | **4.30e-04** |
| Macro F1 Score | **2.66e-02** | **2.62e-03** | **1.12e-02** | **1.18e-03** | **2.55e-06** | **1.23e-04** | **3.14e-03** |
| Micro F1 Score | **1.83e-03** | **1.60e-04** | **1.54e-02** | **3.96e-03** | **2.73e-07** | **7.70e-07** | **4.30e-04** |
| Precision | **1.25e-02** | **9.37e-03** | **3.14e-03** | **6.78e-03** | **1.28e-05** | **1.27e-04** | **1.06e-02** |
| Recall | **4.87e-02** | **1.97e-03** | **2.33e-02** | **1.28e-03** | **3.67e-06** | **7.86e-05** | **6.26e-03** |
| Correlation ($R^2$) | **2.08e-05** | **3.97e-05** | **4.28e-05** | **3.94e-09** | **2.01e-07** | **2.18e-11** | **2.45e-06** |

**Table S3.** The p-values for two-sided t-tests on evaluation metrics for the AR and standard scVI models tested on the Kidney dataset. Bold indicates statistical significance as determined by *P* < 0.05. Results are based on five replicates.

| # Atlas Cells | 0 | 1 | 10 | 100 | 1000 | 10000 | 50000 |
| --- | --- | --- | --- | --- | --- | --- | --- |
| Accuracy | **4.10e-02** | **3.72e-02** | 9.38e-02 | **6.75e-03** | **1.44e-04** | **1.56e-05** | **2.29e-04** |
| Macro F1 Score | **2.11e-02** | **9.16e-03** | **1.18e-02** | **8.74e-03** | **2.40e-05** | **1.35e-05** | **6.56e-04** |
| Micro F1 Score | **4.10e-02** | **3.72e-02** | 9.38e-02 | **6.75e-03** | **1.44e-04** | **1.56e-05** | **2.29e-04** |
| Precision | **1.14e-02** | **2.37e-05** | **2.32e-05** | **1.91e-03** | **5.55e-05** | **3.69e-06** | **2.67e-03** |
| Recall | 5.07e-02 | **2.04e-02** | 7.82e-02 | **1.57e-02** | **2.21e-05** | **1.66e-05** | **2.25e-04** |
| Correlation ($R^2$) | **4.81e-07** | **8.99e-07** | **4.53e-05** | **1.96e-10** | **2.64e-07** | **6.01e-07** | **2.89e-03** |

**Table S4.** The p-values for two-sided t-tests on evaluation metrics for the AR and standard scVI models tested on the Neuron dataset. Bold indicates statistical significance as determined by *P* < 0.05. Results are based on five replicates.

| # Atlas Cells | 0 | 1 | 10 | 100 | 1000 | 10000 | 50000 |
| --- | --- | --- | --- | --- | --- | --- | --- |
| Accuracy | **8.77e-03** | **1.80e-02** | 9.06e-02 | **1.33e-04** | **1.91e-07** | **9.52e-07** | 1.62e-01 |
| Macro F1 Score | **5.91e-03** | **2.33e-03** | 1.72e-01 | **3.48e-04** | **9.48e-07** | **3.99e-06** | 5.76e-01 |
| Micro F1 Score | **8.77e-03** | **1.80e-02** | 9.06e-02 | **1.33e-04** | **1.91e-07** | **9.52e-07** | 1.62e-01 |
| Precision | 7.97e-02 | **3.46e-02** | 1.21e-01 | **1.01e-04** | **3.11e-05** | **6.11e-05** | 8.89e-01 |
| Recall | **6.00e-03** | **2.78e-03** | 1.87e-01 | **2.24e-04** | **5.55e-07** | **1.51e-06** | 5.25e-01 |
| Correlation ($R^2$) | **7.35e-05** | **6.87e-05** | **6.18e-05** | **1.51e-10** | **1.19e-05** | **4.52e-06** | 5.29e-01 |

**Figure S1.**
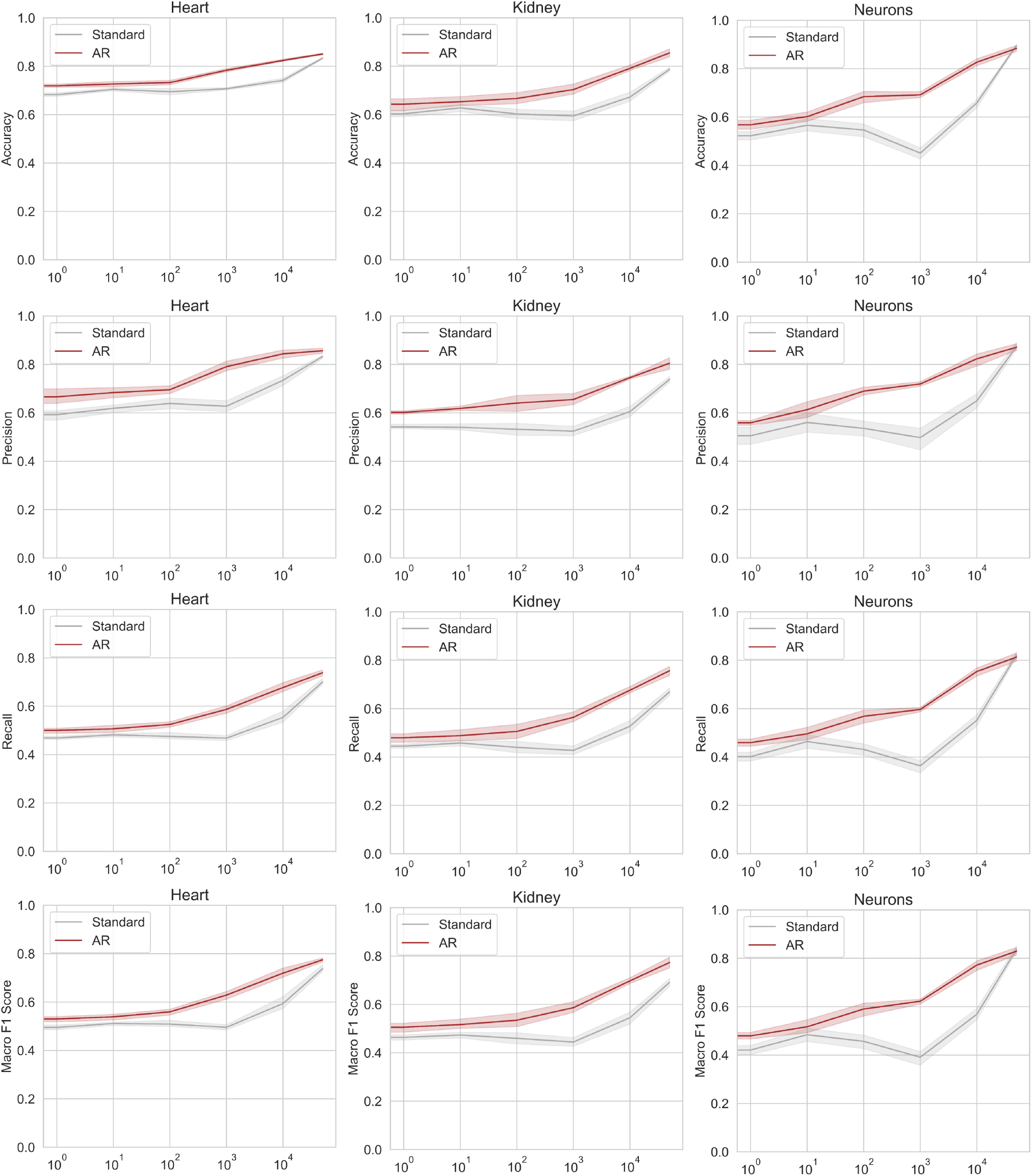
Cell type classification performance on Heart, Kidney, and Neuron evaluation datasets. Classification is done using the model embeddings from scVI models trained either in the standard way (grey) or with AR (red). Performance is measured by the accuracy, precision, recall, and macro F1 scores. The x-axis represents the number of atlas cells included in the blood-based training dataset, with all training sets having a total of 100k cells. Results are based on five different random seeds, with error bars representing the 95% confidence interval across replicates.

**Figure S2.**
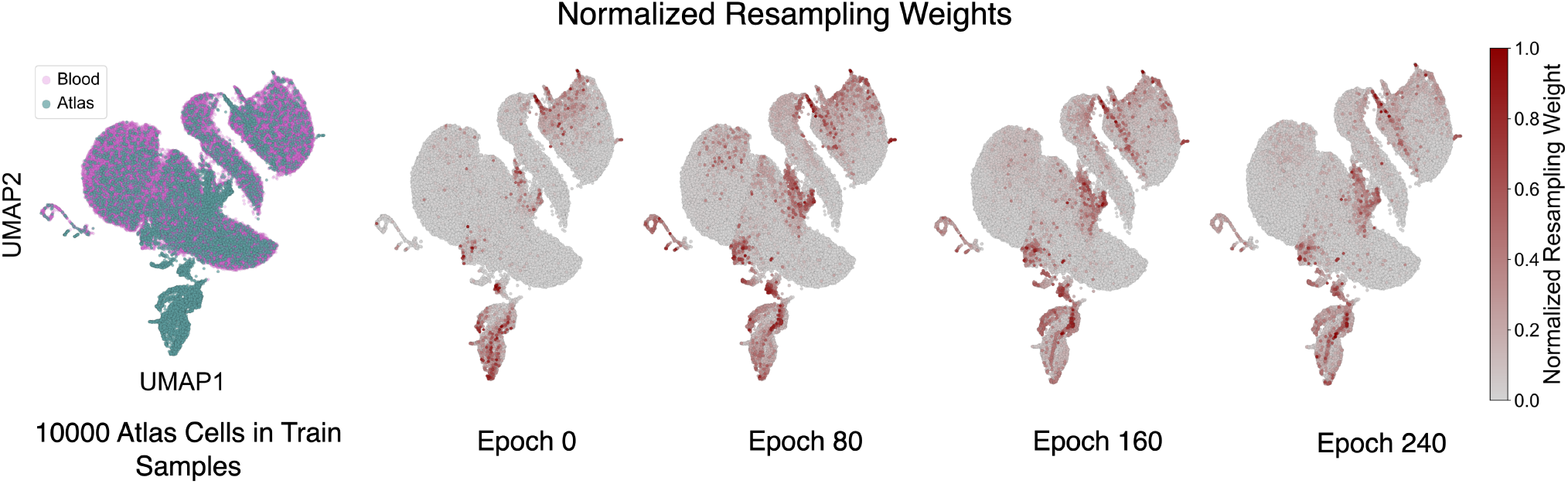
Underrepresented atlas cells are assigned higher resampling weights compared to blood cells in the training corpus. Uniform manifold approximation and projection (UMAP) visualizations of training samples for one random seed of the 90,000 blood cell + 10,000 atlas cell dataset, annotated by their cell group (left; blood in purple and atlas in green) and by normalized resampling weights over the course of AR model training (right).

**Figure S3.**
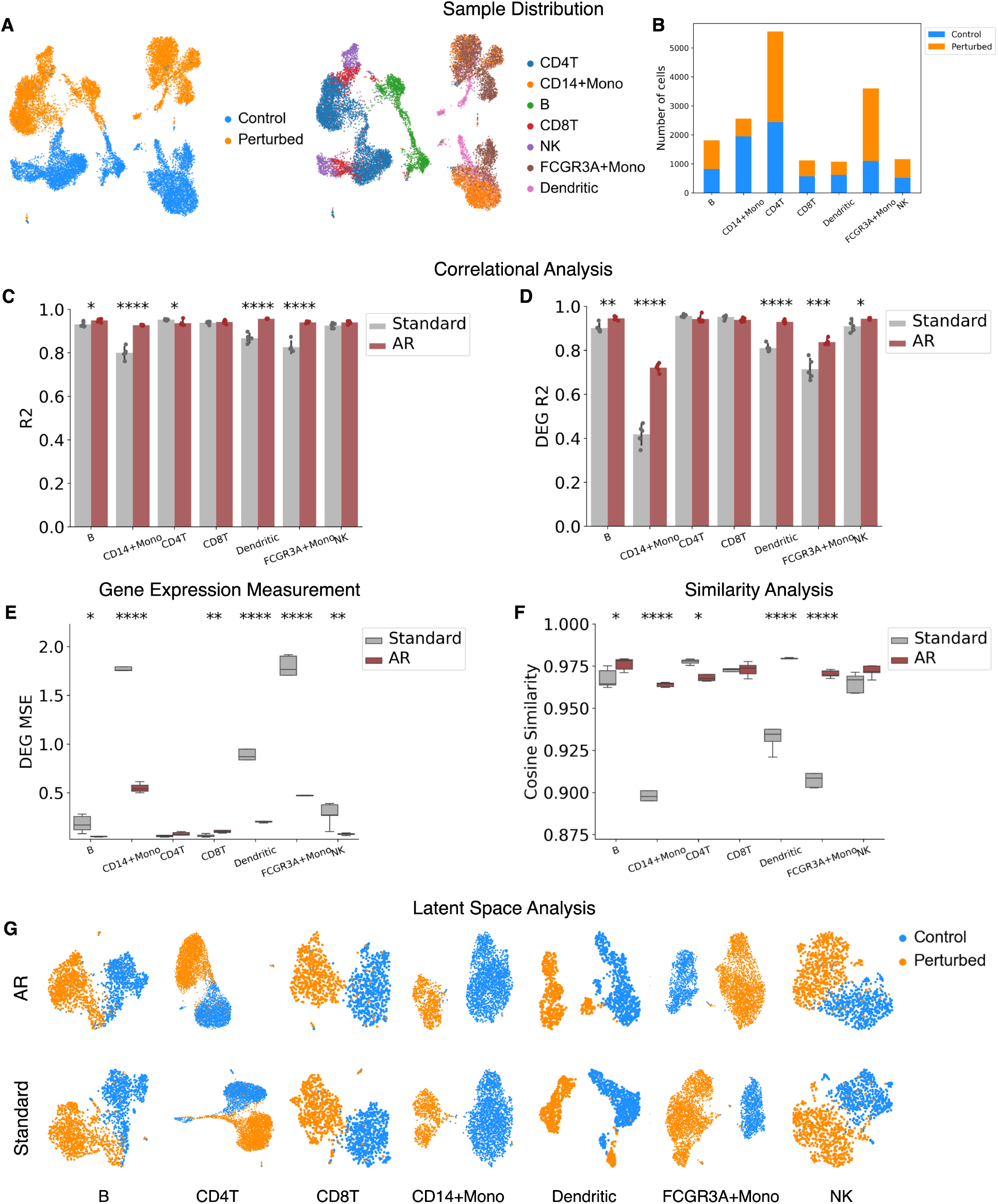
Adaptive resampling enables robust prediction of cellular response to IFN-*β* stimulation in human immune cells. **(A)** UMAP visualization of sample distributions in the PBMC dataset, annotated by condition (control versus perturbed, left) and by cell groups (right). **(B)** Cell counts per condition and cell group. **(C-D)** Square of the Pearson correlation coefficient between real and predicted perturbed gene expression using AR versus standard models for each held-out test cell group across (C) all genes and (D) only differentially expressed genes. Results are based on five different random seeds, with error bars representing the standard deviations across replicates. **(E)** Mean sqaured error (MSE) computed between ground-truth and predicted perturbed gene expression profiles across differentially expressed genes for AR versus standard models. **(F)** Distributions of cosine similarity scores for predicted expression profiles across all genes, relative to ground-truth expression profiles. **(G)** UMAP visualizations of the latent space projections for test samples, annotated by condition, comparing AR and standard learned latent embeddings. Two-sided t-tests were performed in all experiments to compare the means of metrics, with ^*^*P* < 0.05, ^**^*P* < 0.01, ^***^*P* < 0.001, and ^****^*P* < 0.0001.

**Figure S4.**
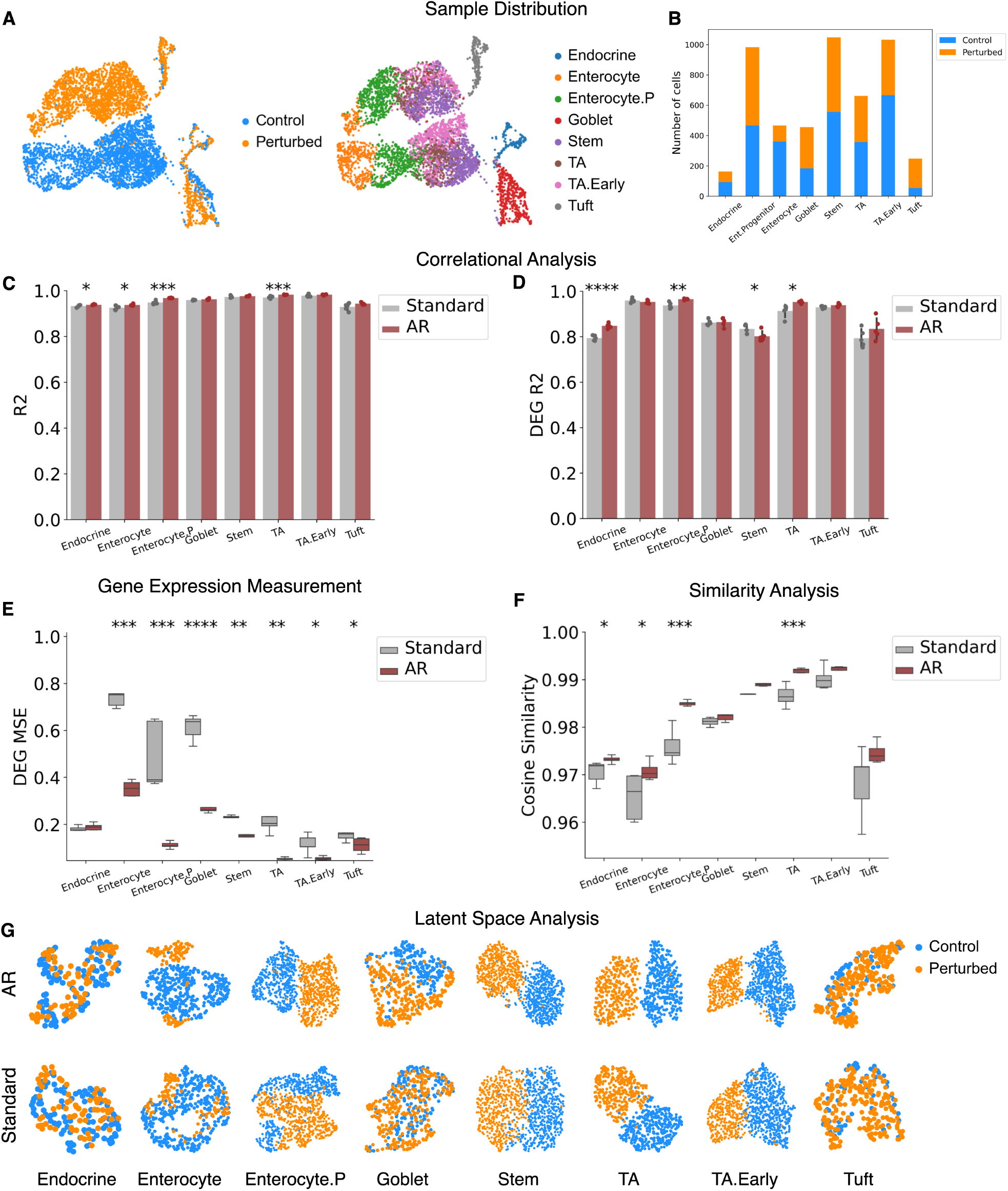
Adaptive resampling enables robust prediction of cellular response to *H. poly* parasitic infection in the murine intestinal epithelium. **(A)** UMAP visualization of sample distributions in the H. poly dataset, annotated by condition (control versus perturbed, left) and by cell groups (right). **(B)** Cell counts per condition and cell group. **(C-D)** Square of the Pearson correlation coefficient between real and predicted perturbed gene expression using AR versus standard models for each held-out test cell group across (C) all genes and (D) only differentially expressed genes. Results are based on five different random seeds, with error bars representing the standard deviations across replicates. **(E)** Mean sqaured error (MSE) computed between ground-truth and predicted perturbed gene expression profiles across differentially expressed genes for AR versus standard models. **(F)** Distributions of cosine similarity scores for predicted expression profiles across all genes, relative to ground-truth expression profiles. **(G)** UMAP visualizations of the latent space projections for test samples, annotated by condition, comparing AR and standard learned latent embeddings. Two-sided t-tests were performed in all experiments to compare the means of metrics, with ^*^*P* < 0.05, ^**^*P* < 0.01, ^***^*P* < 0.001, and ^****^*P* < 0.0001.

**Figure S5.**
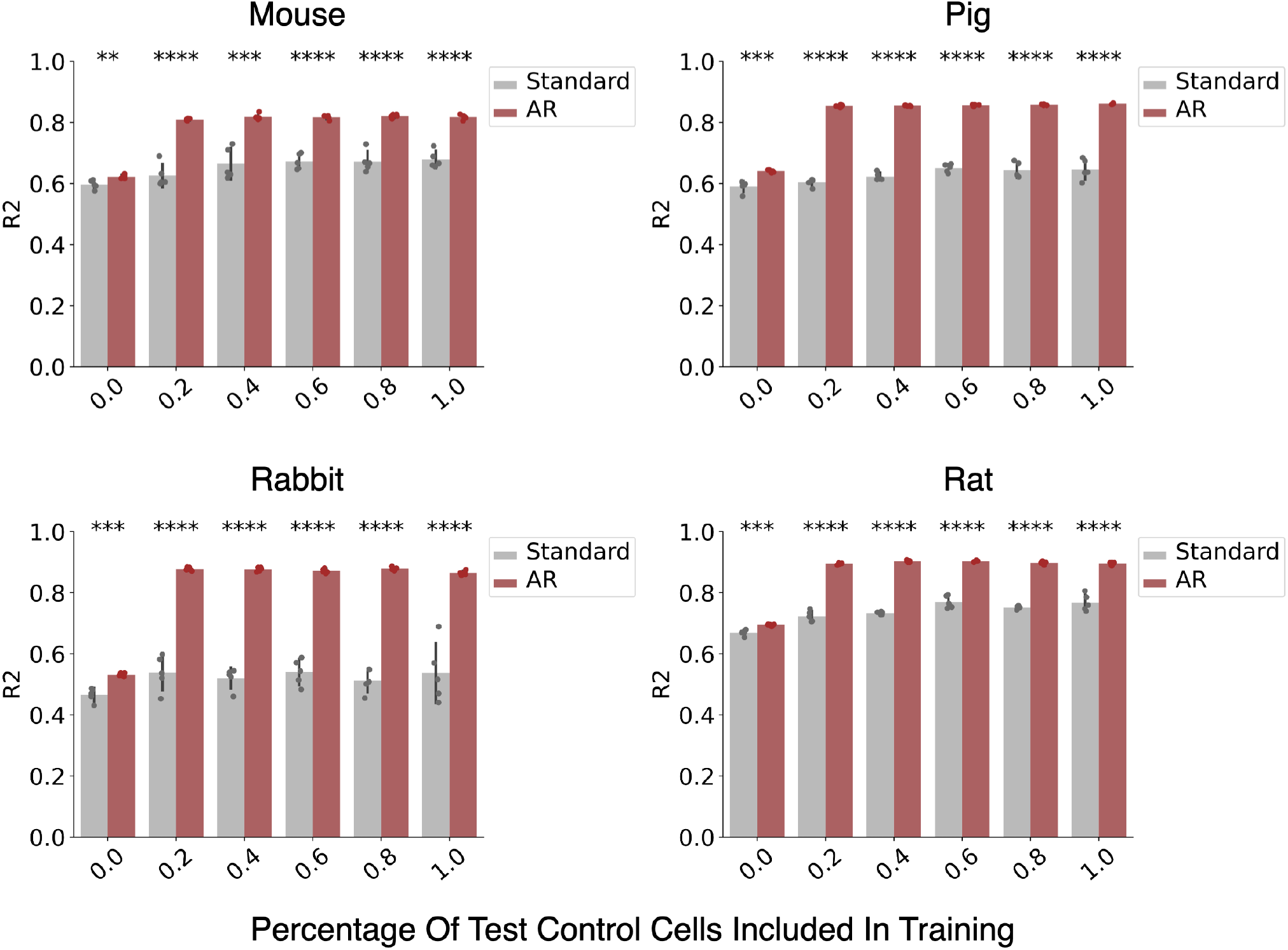
Adaptive resampling enables generalizable perturbation response prediction on species that are underrepresented or completely unseen. Square of the Pearson correlation coefficient (*R*^2^) between ground-truth and predicted perturbed gene expression in rare sample generalization scenarios on the Species dataset for AR versus standard models. Individual plots show results where we trained models while incrementally introducing varying proportions of control cells from the held-out group into the training corpus—specifically, 0%, 20%, 40%, 60%, 80%, and 100%. Results are based on five different random seeds, with error bars representing the standard deviations across replicates. Two-sided t-tests were performed in all experiments to compare the means of metrics, with ^*^*P* < 0.05, ^**^*P* < 0.01, ^***^*P* < 0.001, and ^****^*P* < 0.0001.

**Figure S6.**
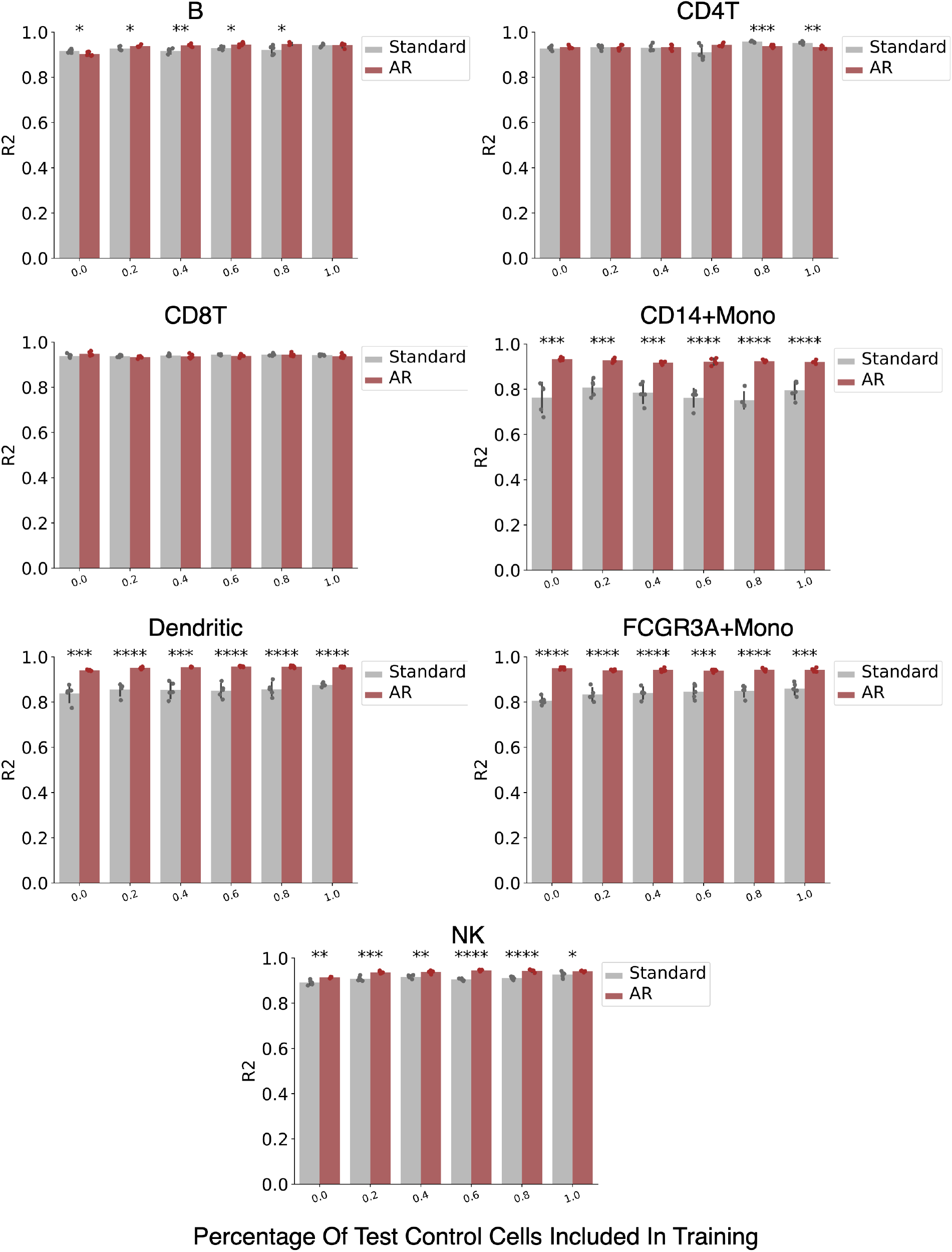
Adaptive resampling enables generalizable perturbation response prediction on underrepresented immune cell populations. Square of the Pearson correlation coefficient (*R*^2^) between ground-truth and predicted perturbed gene expression in rare sample generalization scenarios on the PBMC dataset for AR versus standard models. Individual plots show results where we trained models while incrementally introducing varying proportions of control cells from the held-out group into the training corpus—sp8ecifically, 0%, 20%, 40%, 60%, 80%, and 100%. Results are based on five different random seeds, with error bars representing the standard deviations across replicates. Two-sided t-tests were performed in all experiments to compare the means of metrics, with ^*^*P* < 0.05, ^**^*P* < 0.01, ^***^*P* < 0.001, and ^****^*P* < 0.0001.

**Figure S7.**
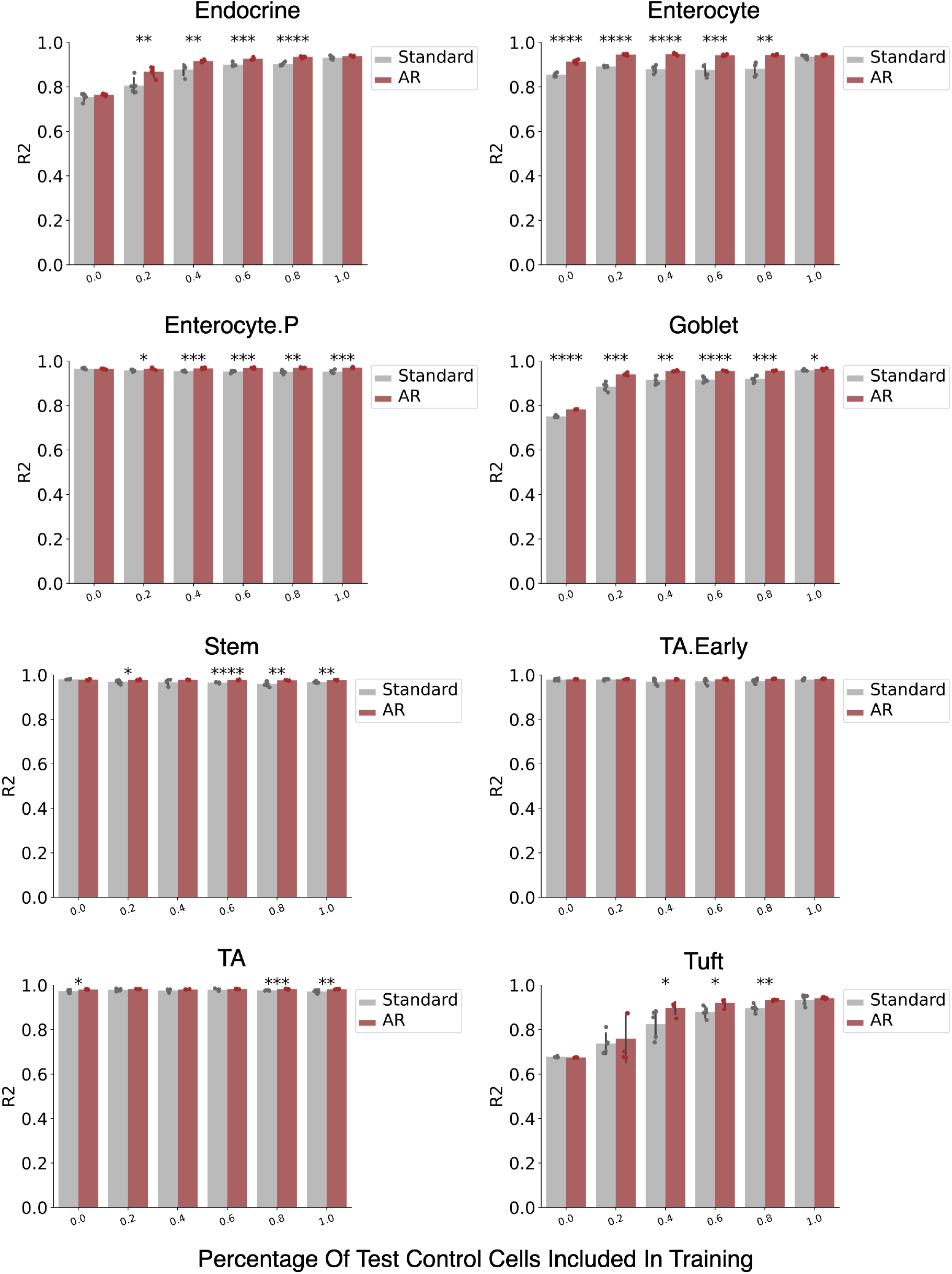
Adaptive resampling enables generalizable perturbation response prediction of parasitic infection in underrepresented intestinal epithelium cells. Square of the Pearson correlation coefficient (*R*^2^) between ground-truth and predicted perturbed gene expression in rare sample generalization scenarios on the H. poly dataset for AR versus standard models. Individual plots show results where we trained models while incrementally introducing varying proportions of control cells from the held-out group into the training corpus—specifically, 0%, 20%, 40%, 60%, 80%, and 100%. Results are based on five different random seeds, with error bars representing the standard deviations across replicates. Two-sided t-tests were performed in all experiments to compare the means of metrics, with ^*^*P* < 0.05, ^**^*P* < 0.01, ^***^*P* < 0.001, and ^****^*P* < 0.0001.

## References

[1] Sonia Nestorowa, Fiona K Hamey, Blanca Pijuan Sala, Evangelia Diamanti, Mairi Shepherd, Elisa Laurenti, Nicola K Wilson, David G Kent, and Berthold Göttgens. A single-cell resolution map of mouse hematopoietic stem and progenitor cell differentiation. Blood, The Journal of the American Society of Hematology, 128(8):e20–e31, 2016.

[2] Lindsey W Plasschaert, Rapolas Žilionis, Rayman Choo-Wing, Virginia Savova, Judith Knehr, Guglielmo Roma, Allon M Klein, and Aron B Jaffe. A single-cell atlas of the airway epithelium reveals the cftr-rich pulmonary ionocyte. Nature, 560(7718):377–381, 2018.

[3] Aashi Jindal, Prashant Gupta, Jayadeva, and Debarka Sengupta. Discovery of rare cells from voluminous single cell expression data. Nature Communications, 9(1):4719, 2018.

[4] Aleksandrina Goeva, Michael-John Dolan, Judy Luu, Eric Garcia, Rebecca Boiarsky, Rajat M Gupta, and Evan Macosko. Hidden: a machine learning method for detection of disease-relevant populations in case-control single-cell transcriptomics data. Nature Communications, 15(1):9468, 2024.

[5] Dragomirka Jovic, Xue Liang, Hua Zeng, Lin Lin, Fengping Xu, and Yonglun Luo. Singlecell rna sequencing technologies and applications: A brief overview. Clinical and Translational Medicine, 12(3):e694, 2022.

[6] Guangzhe Ge, Yang Han, Jianye Zhang, Xinxin Li, Xiaodan Liu, Yanqing Gong, Zhentao Lei, Jie Wang, Weijie Zhu, Yangyang Xu, et al. Single-cell rna-seq reveals a developmental hierarchy super-imposed over subclonal evolution in the cellular ecosystem of prostate cancer. Advanced Science, 9(15):2105530, 2022.

[7] Itay Tirosh, Andrew S Venteicher, Christine Hebert, Leah E Escalante, Anoop P Patel, Keren Yizhak, Jonathan M Fisher, Christopher Rodman, Christopher Mount, Mariella G Filbin, et al. Single-cell rna-seq supports a developmental hierarchy in human oligodendroglioma. Nature, 539(7628):309–313, 2016.

[8] Lu Pan, Paolo Parini, Roman Tremmel, Joseph Loscalzo, Volker M Lauschke, Bradley A Maron, Paola Paci, Ingemar Ernberg, Nguan Soon Tan, Zehuan Liao, et al. Single cell atlas: a single-cell multi-omics human cell encyclopedia. Genome Biology, 25(1):104, 2024.

[9] Jingyao Zeng, Yadong Zhang, Yunfei Shang, Jialin Mai, Shuo Shi, Mingming Lu, Congfan Bu, Zhewen Zhang, Zaichao Zhang, Yang Li, et al. CancerSCEM: a database of single-cell expression map across various human cancers. Nucleic Acids Research, 50(D1):D1147– D1155, 2022.

[10] Jesse Zhang, Airol A Ubas, Richard de Borja, Valentine Svensson, Nicole Thomas, Neha Thakar, Ian Lai, Aidan Winters, Umair Khan, Matthew G Jones, et al. Tahoe-100m: A gigascale single-cell perturbation atlas for context-dependent gene function and cellular modeling. BioRxiv, pages 2025–02, 2025.

[11] Malte D Luecken and Fabian J Theis. Current best practices in single-cell rna-seq analysis: a tutorial. Molecular Systems Biology, 15(6):e8746, 2019.

[12] Haotian Cui, Chloe Wang, Hassaan Maan, Kuan Pang, Fengning Luo, Nan Duan, and Bo Wang. scgpt: toward building a foundation model for single-cell multi-omics using generative ai. Nature Methods, pages 1–11, 2024.

[13] Fan Yang, Wenchuan Wang, Fang Wang, Yuan Fang, Duyu Tang, Junzhou Huang, Hui Lu, and Jianhua Yao. scbert as a large-scale pretrained deep language model for cell type annotation of single-cell rna-seq data. Nature Machine Intelligence, 4(10):852–866, 2022.

[14] Wenpin Hou and Zhicheng Ji. Assessing GPT-4 for cell type annotation in single-cell RNA-seq analysis. Nature Methods, pages 1–4, 2024.

[15] Artur Szałata, Karin Hrovatin, Sören Becker, Alejandro Tejada-Lapuerta, Haotian Cui, Bo Wang, and Fabian J Theis. Transformers in single-cell omics: a review and new perspectives. Nature Methods, 21(8):1430–1443, 2024.

[16] Christina V Theodoris, Ling Xiao, Anant Chopra, Mark D Chaffin, Zeina R Al Sayed, Matthew C Hill, Helene Mantineo, Elizabeth M Brydon, Zexian Zeng, X Shirley Liu, et al. Transfer learning enables predictions in network biology. Nature, 618(7965):616–624, 2023.

[17] Theresa Willem, Vladimir A Shitov, Malte D Luecken, Niki Kilbertus, Stefan Bauer, Marie Piraud, Alena Buyx, and Fabian J Theis. Biases in machine-learning models of human singlecell data. Nature Cell Biology, pages 1–9, 2025.

[18] Ajay Nadig, Akshaya Thoutam, Madeline Hughes, Anay Gupta, Andrew W Navia, Nicolo Fusi, Srivatsan Raghavan, Peter S Winter, Ava P Amini, and Lorin Crawford. Consequences of training data composition for deep learning models in single-cell biology. bioRxiv, pages 2025–02, 2025.

[19] Alan DenAdel, Madeline Hughes, Akshaya Thoutam, Anay Gupta, Andrew W Navia, Nicolo Fusi, Srivatsan Raghavan, Peter S Winter, Ava P Amini, and Lorin Crawford. Evaluating the role of pre-training dataset size and diversity on single-cell foundation model performance. bioRxiv, pages 2024–12, 2024.

[20] Alexander Amini, Ava P Soleimany, Wilko Schwarting, Sangeeta N Bhatia, and Daniela Rus. Uncovering and mitigating algorithmic bias through learned latent structure. In Proceedings of the 2019 AAAI/ACM Conference on AI, Ethics, and Society, pages 289–295, 2019.

[21] Hung S Luu. Laboratory data as a potential source of bias in healthcare artificial intelligence and machine learning models. Annals of Laboratory Medicine, 45(1):12–21, 2025.

[22] Atta Ur Rahman, Bibi Saqia, Yousef S Alsenani, and Inam Ullah. Data quality, bias, and strategic challenges in reinforcement learning for healthcare: A survey. International Journal of Data Informatics and Intelligent Computing, 3(3):24–42, 2024.

[23] Jiashuo Liu, Zheyan Shen, Yue He, Xingxuan Zhang, Renzhe Xu, Han Yu, and Peng Cui. Towards out-of-distribution generalization: A survey. arXiv preprint 2108.13624, 2021.

[24] Tianhao Li, Zixuan Wang, Yuhang Liu, Sihan He, Quan Zou, and Yongqing Zhang. An overview of computational methods in single-cell transcriptomic cell type annotation. Briefings in Bioinformatics, 26(3), 2025.

[25] Shudong Wang, Hengxiao Li, Kuijie Zhang, Hao Wu, Shanchen Pang, Wenhao Wu, Lan Ye, Jionglong Su, and Yulin Zhang. scsid: A lightweight algorithm for identifying rare cell types by capturing differential expression from single-cell sequencing data. Computational and Structural Biotechnology Journal, 23:589–600, 2024.

[26] Mario L Suvà and Itay Tirosh. Single-cell rna sequencing in cancer: lessons learned and emerging challenges. Molecular Cell, 75(1):7–12, 2019.

[27] Zilong Zhang, Feifei Cui, Chen Lin, Lingling Zhao, Chunyu Wang, and Quan Zou. Critical downstream analysis steps for single-cell rna sequencing data. Briefings in Bioinformatics, 22(5):bbab105, 2021.

[28] Sumeer Ahmad Khan, Alberto Maillo, Vincenzo Lagani, Robert Lehmann, Narsis A Kiani, David Gomez-Cabrero, and Jesper Tegner. Reusability report: Learning the transcriptional grammar in single-cell rna-sequencing data using transformers. Nature Machine Intelligence, pages 1–10, 2023.

[29] Hassaan Maan, Lin Zhang, Chengxin Yu, Michael Geuenich, Kieran R Campbell, and Bo Wang. The differential impacts of dataset imbalance in single-cell data integration. BioRxiv, pages 2022–10, 2022.

[30] Abdel Rahman Alsabbagh, Albert Maillo Ruiz de Infante, David Gomez-Cabrero, Narsis Kiani, Sumeer Ahmad Khan, and Jesper N Tegner. Foundation models meet imbalanced single-cell data when learning cell type annotations. BioRxiv, pages 2023–10, 2023.

[31] Seungbyn Baek, Kyungwoo Song, and Insuk Lee. Single-cell foundation models: bringing artificial intelligence into cell biology. Experimental & Molecular Medicine, pages 1–13, 2025.

[32] Charlotte Bunne, Yusuf Roohani, Yanay Rosen, Ankit Gupta, Xikun Zhang, Marcel Roed, Theo Alexandrov, Mohammed AlQuraishi, Patricia Brennan, Daniel B Burkhardt, et al. How to build the virtual cell with artificial intelligence: priorities and opportunities. Cell, 187(25): 7045–7063, 2024.

[33] Yuansong Zeng, Jiancong Xie, Ningyuan Shangguan, Zhuoyi Wei, Wenbing Li, Yun Su, Shuangyu Yang, Chengyang Zhang, Jinbo Zhang, Nan Fang, et al. CellFM: a large-scale foundation model pre-trained on transcriptomics of 100 million human cells. Nature Communications, 16(1):4679, 2025.

[34] Chloe Wang, Haotian Cui, Andrew Zhang, Ronald Xie, Hani Goodarzi, and Bo Wang. scGPT-spatial: Continual pretraining of single-cell foundation model for spatial transcriptomics, 2025.

[35] Michael J Geuenich, Dae-won Gong, and Kieran R Campbell. The impacts of active and self-supervised learning on efficient annotation of single-cell expression data. Nature Communications, 15(1):1014, 2024.

[36] Jian Jiang, Chunhuan Zhang, Lu Ke, Nicole Hayes, Yueying Zhu, Huahai Qiu, Bengong Zhang, Tianshou Zhou, and Guo-Wei Wei. A review of machine learning methods for imbalanced data challenges in chemistry. Chemical Science, 2025.

[37] Yuqi Cheng, Xingyu Fan, Jianing Zhang, and Yu Li. A scalable sparse neural network framework for rare cell type annotation of single-cell transcriptome data. Communications Biology, 6(1):545, 2023.

[38] Changde Cheng, Wenan Chen, Hongjian Jin, and Xiang Chen. A review of single-cell RNAseq annotation, integration, and cell–cell communication. Cells, 12(15):1970, 2023.

[39] Linh Truong, Thao Truong, and Huy Nguyen. Automated cell annotation in scRNA-seq data using unique marker gene sets. bioRxiv, 2024.

[40] Romain Lopez, Jeffrey Regier, Michael B Cole, Michael I Jordan, and Nir Yosef. Deep generative modeling for single-cell transcriptomics. Nature Methods, 15(12):1053–1058, 2018.

[41] Mohammad Lotfollahi, F Alexander Wolf, and Fabian J Theis. scGen predicts single-cell perturbation responses. Nature Methods, 16(8):715–721, 2019.

[42] Ihab Bendidi, Shawn Whitfield, Kian Kenyon-Dean, Hanene Ben Yedder, Yassir El Mesbahi, Emmanuel Noutahi, and Alisandra K Denton. Benchmarking transcriptomics foundation models for perturbation analysis: one PCA still rules them all. arXiv preprint 2410.13956, 2024.

[43] Kasia Z Kedzierska, Lorin Crawford, Ava P Amini, and Alex X Lu. Zero-shot evaluation reveals limitations of single-cell foundation models. Genome Biology, 26(1):101, 2025.

[44] Felix Fischer, David S. Fischer, Roman Mukhin, Andrey Isaev, Evan Biederstedt, Alexandra-Chloé Villani and Fabian J. Theis. scTab: Scaling cross-tissue single-cell annotation models. Nature Communications, 15(1):6611, Aug 2024.

[45] Tzachi Hagai, Xi Chen, Ricardo J Miragaia, Raghd Rostom, Tomás Gomes, Natalia Kunowska, Johan Henriksson, Jong-Eun Park, Valentina Proserpio, Giacomo Donati, et al. Gene expression variability across cells and species shapes innate immunity. Nature, 563 (7730):197–202, 2018.

[46] Hyun Min Kang, Meena Subramaniam, Sasha Targ, Michelle Nguyen, Lenka Maliskova, Elizabeth McCarthy, Eunice Wan, Simon Wong, Lauren Byrnes, Cristina M Lanata, et al. Multiplexed droplet single-cell RNA-sequencing using natural genetic variation. Nature Biotechnology, 36(1):89–94, 2018.

[47] Adam L Haber, Moshe Biton, Noga Rogel, Rebecca H Herbst, Karthik Shekhar, Christopher Smillie, Grace Burgin, Toni M Delorey, Michael R Howitt, Yarden Katz, et al. A single-cell survey of the small intestinal epithelium. Nature, 551(7680):333–339, 2017.

[48] Constantin Ahlmann-Eltze, Wolfgang Huber, and Simon Anders. Deep-learning-based gene perturbation effect prediction does not yet outperform simple linear baselines. Nature Methods, pages 1–5, 2025.

[49] Gerold Csendes, Gema Sanz, Kristóf Z Szalay, and Bence Szalai. Benchmarking foundation cell models for post-perturbation RNA-seq prediction. BMC Genomics, 26(1):393, 2025.

[50] Wei Lan, Guohang He, Mingyang Liu, Qingfeng Chen, Junyue Cao, and Wei Peng. Transformer-based single-cell language model: A survey. Big Data Mining and Analytics, 7 (4):1169–1186, 2024.

[51] Nicholas D Youngblut, Christopher Carpenter, Jaanak Prashar, Chiara Ricci-Tam, Rajesh Ilango, Noam Teyssier, Silvana Konermann, Patrick D Hsu, Alexander Dobin, David P Burke, et al. scBaseCount: an ai agent-curated, uniformly processed, and continually expanding single cell data repository. bioRxiv, 2025.

[52] Chanseok Kang. Debiasing facial detection systems. https://goodboychan.github.io/python/tensorflow/mit/2021/02/27/Debiasing.html, 2021.

[53] Adam Gayoso, Romain Lopez, Galen Xing, Pierre Boyeau, Valeh Valiollah Pour Amiri, Justin Hong, Katherine Wu, Michael Jayasuriya, Edouard Mehlman, Maxime Langevin, et al. A python library for probabilistic analysis of single-cell omics data. Nature Biotechnology, 40(2):163–166, 2022.

[54] CZI Cell Science Program, Shibla Abdulla, Brian Aevermann, Pedro Assis, Seve Badajoz, Sidney M Bell, Emanuele Bezzi, Batuhan Cakir, Jim Chaffer, Signe Chambers, et al. CZ CELLxGENE discover: a single-cell data platform for scalable exploration, analysis and modeling of aggregated data. Nucleic Acids Research, 53(D1):D886–D900, 2025.

[55] Kimberly Siletti, Rebecca Hodge, Alejandro Mossi Albiach, Ka Wai Lee, Song-Lin Ding, Lijuan Hu, Peter Lönnerberg, Trygve Bakken, Tamara Casper, Michael Clark, Nick Dee, Jessica Gloe, Daniel Hirschstein, Nadiya V Shapovalova, C Dirk Keene, Julie Nyhus, Herman Tung, Anna Marie Yanny, Ernest Arenas, Ed S Lein, and Sten Linnarsson. Transcriptomic diversity of cell types across the adult human brain. Science, 382(6667):eadd7046, oct 2023.

